# Synapse-type-specific competitive Hebbian learning forms functional recurrent networks

**DOI:** 10.1101/2022.03.11.483899

**Authors:** Samuel Eckmann, Edward James Young, Julijana Gjorgjieva

## Abstract

Cortical networks exhibit complex stimulus-response patterns that are based on specific recurrent interactions between neurons. For example, the balance between excitatory and inhibitory currents has been identified as a central component of cortical computations. However, it remains unclear how the required synaptic connectivity can emerge in developing circuits where synapses between excitatory and inhibitory neurons are simultaneously plastic. Using theory and modeling, we propose that a wide range of cortical response properties can arise from a single plasticity paradigm that acts simultaneously at all excitatory and inhibitory connections – Hebbian learning that is stabilized by the synapse-type-specific competition for a limited supply of synaptic resources. In plastic recurrent circuits, this competition enables the formation and decorrelation of inhibition-balanced receptive fields. Networks develop an assembly structure with stronger synaptic connections between similarly tuned excitatory and inhibitory neurons and exhibit response normalization and orientation-specific center-surround suppression, reflecting the stimulus statistics during training. These results demonstrate how neurons can self-organize into functional networks and suggest an essential role for synapse-type-specific competitive learning in the development of cortical circuits.

**Significance Statement:** Cortical circuits perform diverse computations, primarily determined by highly structured synaptic connectivity patterns that develop during early sensory experience via synaptic plasticity. To understand how these structured connectivity patterns emerge, we introduce a general learning framework for networks of recurrently connected neurons. The framework is rooted in the biologically plausible assumption that synapses compete for limited synaptic resources, which stabilizes synaptic growth. Motivated by the unique protein composition of different synapse types, we assume that different synapse types compete for separate resource pools. Using theory and simulation, we show how this synapse-type-specific competition allows the stable development of structured synaptic connectivity patterns, as well as diverse computations like response normalization and surround suppression.

---

Computation in neural circuits is based on the interactions between recurrently connected excitatory (E) and inhibitory (I) neurons (*1*–*4*). In sensory cortices, response normalization, surround and gain modulation, predictive processing, and attention all critically involve inhibitory neurons (*5*– *10*). Theoretical work has highlighted the experimentally observed balance of stimulus selective excitatory and inhibitory input currents as a critical requirement for many neural computations (*11*–*16*). For example, recent models based on balanced E-I networks can explain a wide range of cortical phenomena, such as cross-orientation and surround suppression (*17, 18*), as well as stimulus-induced neural variability (*19*–*21*). A major caveat of these models is that the network connectivity is usually static and designed by hand, albeit based on experimental measurements. In contrast, in the brain, synapses are plastic and adjust to the statistics of sensory inputs. How synaptic weights self-organize in a biologically plausible manner to generate many of the non-linear response properties observed experimentally is not well understood. Earlier theoretical work on inhibitory plasticity has focused on the balance of excitation and inhibition in single neurons (*22*–*24*), but has not been able to explain the development of inhibition-balanced receptive fields when excitatory and inhibitory inputs are both plastic. In more recent recurrent network models, only a fraction of excitatory and inhibitory synapse-types are modeled as plastic and neural responses exhibit a narrow subset of the different response patterns recorded in experiments (*14, 25*–*29*).

Here we present a Hebbian learning framework with minimal assumptions that explains a wide range of experimental observations. Our framework is based on two key properties: First, all synaptic strengths evolve according to a Hebbian plasticity rule that is stabilized by the competition for a limited supply of synaptic resources (*30*–*33*). Second, motivated by the unique protein composition of excitatory and inhibitory synapses, different synapse-types compete for separate resource pools. Building on classical work on Hebbian plasticity (*30, 31*), we develop an analytical framework that provides an intuitive understanding of the weight dynamics in recurrent networks of excitatory and inhibitory neurons. In numerical simulations, we reveal how the synapse-type-specific competition for resources enables the self-organization of neurons into functional networks. Beyond the formation of inhibition-balanced feedforward receptive fields, we demonstrate that emergent recurrent connectivity can generate a wide range of computations observed in cortical circuits.

## Results

### Synapse-type-specific plasticity enables the joint development of stimulus selectivity and E-I balance

To understand plasticity in recurrently connected E-I networks, we considered simplified circuits of increasing complexity. We first asked how E-I balance and stimulus selectivity can simultaneously develop in a single neuron. The neuron receives input from an upstream population of excitatory neurons, and disynaptic inhibitory input from a population of laterally connected inhibitory neurons that themselves receive input from the same upstream population (Fig. 1*A*). We studied the self-organization of excitatory and inhibitory synapses that project onto the single postsynaptic neuron (Fig. 1*B*), assuming that input synapses that project onto inhibitory neurons remained fixed (Fig. 1*A*). Following experimental results (*34*–*37*), we assumed that inhibitory and excitatory input neurons are equally selective for the orientation of a stimulus grating (Fig. 1*C*, bottom). We presented uniformly distributed oriented stimuli to the network in random order. Stimuli elicited a Gaussian-shaped response in the population of input neurons (Fig. 1*C*, top) and thus drove the postsynaptic neuron (see Methods for details). Synapses are plastic according to a basic Hebbian rule:

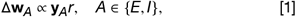

where *r* is the postsynaptic firing rate, **y**_*A*_ is a vector that holds the presynaptic firing rates of excitatory (*A* = *E*) and inhibitory (*A* = *I*) neurons, and Δ**w**_*A*_ are the corresponding synaptic weight changes. Experimental results have shown that after the induction of long-term plasticity neither the total excitatory nor the total inhibitory synaptic area change (*32*). This suggests that a synapse can only grow at the expense of another synapse — a competitive mechanism potentially mediated by the limited supply of synaptic proteins (Fig. 1*D*) (*33*). Motivated by these results, we adopted a competitive normalization rule for both excitatory and inhibitory synapses:

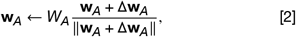

where *A* ∈ {*E, I* }, and *W*_*E*_, *W*_*I*_ are the maintained total excitatory and inhibitory synaptic weight, respectively. Shortly after random initialization, excitatory and inhibitory weights stabilize (Fig. 1*E*) and form aligned, Gaussian-shaped tuning curves (Fig. 1*F*) that reflect the shape of the input stimuli (Fig. 1*C*). As a result, neural responses become orientation selective while inhibitory and excitatory inputs are equally tuned, which demon-strates the joint development of stimulus selectivity and E-I balance.

**Figure 1:**
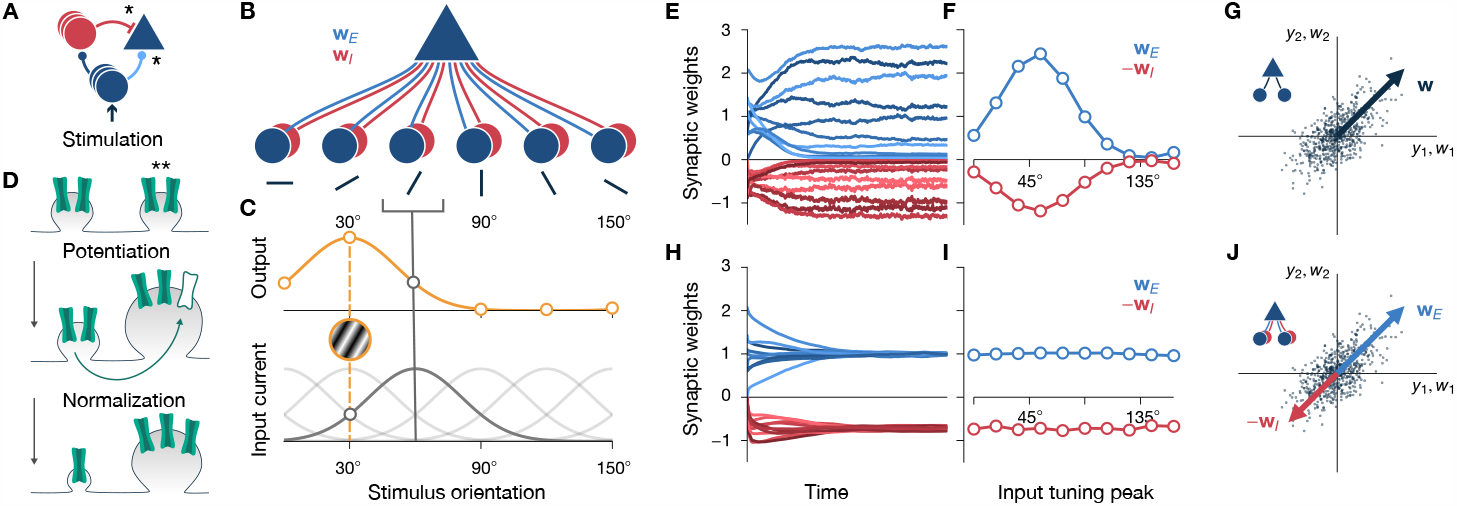
Synapse-type-specific competitive Hebbian learning enables the development of stimulus selectivity and inhibitory balance. (*A*) Feedforward input to a model pyramidal neuron during (blue triangle) stimulation. The neuron receives direct excitation (lightblue) and disynaptic inhibition (red). Plastic synapses are marked by *. (*B*) A single postsynaptic pyramidal neuron receives synaptic input from a population of excitatory (**w**_*E*_), and a population of inhibitory (**w**_*I*_) neurons. (*C*) Excitatory and inhibitory input neurons are equally tuned to the orientation of a stimulus grating (bottom, tuning curve of neurons tuned to 60° highlighted in dark gray) and exhibit a Gaussian-shaped population response (orange, solid line) when a single grating of 30° is presented (orange plate, dashed line). (*D*) Hebbian potentiation of a synapse (**) is normalized due to a limited amount of synaptic resources in the dendritic branch, here reflected by a fixed number of synaptic channels (green). (*E*) Weight convergence of synapses of the feedforward circuit in *B*, where excitatory (blue) and inhibitory (red) weights are plastic according to synapse-type-specific competitive Hebbian learning. All synaptic weights were initialized randomly. (*F*) Final synaptic weight strengths, after training, as a function of the tuning peak of the corresponding presynaptic neurons. (*G*) Excitatory synaptic weight vector (blue arrow) of a single pyramidal neuron with linear activation function. The pyramidal neuron receives input from two excitatory neurons (*y*_1_ and *y*_2_, compare inset). Each dot corresponds to one input pattern. After training, the weight vector aligns with the direction of maximum variance, which corresponds to the principal eigenvector of the input covariance matrix. (*H & I*) Same as in *E* and *F*, but for classic inhibitory plasticity. The development of stimulus selectivity is prevented by fast inhibitory plasticity. (*J*) Excitatory (blue) and inhibitory (red) synaptic weight vectors of a single pyramidal neuron with linear activation function. The pyramidal neuron receives input from two pairs of excitatory and inhibitory neurons (*y*_1_ and *y*_2_, compare inset). Each excitatory-inhibitory input pair has identical firing activities *y*_*i*_. After training via synapse-type-specific competitive Hebbian learning, the excitatory and inhibitory weight vectors both align with the principal component, i.e., excitatory and inhibitory synaptic weights are balanced.

### Excitatory plasticity performs principal component analysis

To uncover the principles of synapse-type-specific competitive Hebbian learning, we analyzed the feedforward model analytically. It is well established that in the absence of inhibition, competitive Hebbian learning rules generate stimulus selective excitatory receptive fields (*30, 31*). In the case of a linear activation function, *r* ∝ *u* ≡ **w**^⊤^**y**, the expected total synaptic efficacy changes can be expressed as (*31*):

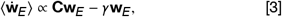

were **C** =⟨ **y**_*E*_ **y**_*E*_^⊤^⟩ is the input covariance matrix, with⟨ ·⟩being the temporal average, and *γ* is a scalar normalization factor that regulates Hebbian growth. Then, fixed points, for which 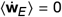, are eigenvectors of the covariance matrix. The neuron becomes selective to the first principal component of its input data, i.e., the fixed point input weight vector aligns with the input space direction of maximum variance (*30, 31*) (Fig. 1*G*; see Supplementary Material (SM) Sec. 1.2 for details). For non-linear activation functions *r* = *f* (*u*), neurons become selective for higher-order correlations, e.g., independent components, in their inputs (*38, 39*). Such learning rules have been shown to result in feedforward receptive fields that resemble simple cell receptive fields in visual cortex (*40, 41*). In the following, we call the fixed points of such pure feedforward circuits ‘input modes’. This entails principal components, in the case of linear activation functions, and more complex, e.g., simple-cell-like, receptive fields in the case of non-linear activation functions.

### Classic inhibitory plasticity prevents stimulus selectivity

We next examined how inhibitory plasticity affects the development of stimulus selectivity. Previous work has suggested that inhibitory synaptic plasticity in the cortex is Hebbian (*42, 43*) and imposes a target firing rate *r*_0_ on the postsynaptic neuron (*23*):

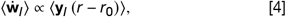

where synaptic change becomes zero when the postsynaptic firing rate *r* is equal to the target rate *r*_0_. With this ‘classic’ inhibitory plasticity rule, inhibitory synaptic weight growth is unbounded. However, since an increase of inhibitory synaptic weights usually entails a decrease in postsynaptic firing rate *r* the plasticity rule is self-limiting and synaptic weights stop growing once the target firing rate *r*_0_ is reached. When excitatory synaptic weights remain fixed, classic inhibitory plasticity leads to balanced excitatory and inhibitory input currents (*23*). However, when excitatory synaptic weights are also plastic, neurons develop no stimulus selectivity (*24*): Classic inhibitory plasticity must act on a faster timescale than excitatory plasticity to maintain stability (*24*). Then the postsynaptic target firing rate is consistently met and average excitatory synaptic weight changes only differ amongst each other due to different average presynaptic firing rates, which prevents the development of stimulus selectivity (Fig. 1*H & I*; see SM Sec. 1.2.3 for details).

### Synapse-type-specific competition enables balanced principal component analysis

Synapse-type-specific competitive Hebbian learning (Eq. 1, and 2) enables the joint development of stimulus selectivity and balanced input currents. In contrast to classic inhibitory plasticity, under synapse-type-specific competitive Hebbian learning, inhibitory synaptic growth is not stabilized by a target firing rate. Instead, as excitatory synapses, inhibitory synapses compete for a limited supply of synaptic resources that maintain the total amount of synaptic strength. As we did for excitatory synapses (Eq. 3), we incorporated the normalization step (Eq. 2) into the update rule (Eq. 1) and considered the simpler case of a linear activation function *f* (*u*) ∝ *u*:

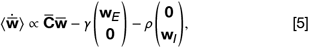

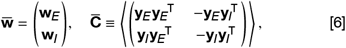

where *γ* and *ρ* are scalars that ensure normalization of excitatory and inhibitory weights, respectively. In addition, we defined the modified covariance matrix 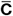. Then multiples of the excitatory and the inhibitory part of the eigenvectors of the modified covariance matrix 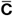 are fixed points of the weight dynamics (see SM Sec. 2 for details). When excitatory and inhibitory inputs are equally stimulus selective, such that one can approximate **y**_*E*_ ∝ **y**_*I*_, the modified covariance matrix 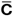 is composed of multiples of the original covariance matrix **C** (cf. Eq. 6). This implies that, if excitatory and inhibitory synaptic weights have identical shape, **w**_*E*_ ∝ **w**_*I*_, equal to a multiple of an eigenvector of **C**, the system is in a fixed point (Fig. 1*J*), where 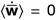 (cf. Eq. 5). Neurons become selective for activity along one particular input direction, while excitatory and inhibitory neural inputs are co-tuned, which explains the joint development of stimulus selectivity and E-I balance in feedforward circuits, in agreement with our numerical simulations with non-linear activation functions (Fig. 1*E & F*).

### Lateral inputs shape feedforward weight dynamics

We wanted to understand how fully plastic recurrent networks of excitatory and inhibitory neurons can self-organize into functional circuits. Therefore, we next investigated the effect of synapse-type-specific competitive Hebbian learning in recurrent networks.

In a first step, we considered how lateral input from an excitatory neuron with fixed selectivity for a specific feedforward input mode affects synaptic weight dynamics in a simple microcircuit motif (Fig. 2*A*, top). We observed that a downstream neuron becomes preferentially tuned to the feedforward input mode of the lateral projecting neuron (Fig. 2*A*, bottom; cf. SM Sec. 3). Similarly, laterally projecting inhibitory neurons repel downstream neurons from their input modes (Fig. 2*B*). However, when two excitatory neurons are reciprocally connected, they pull each other towards their respective input modes, and their tuning curves and activities become correlated (Fig. 2*C*). This contradicts experimental observations that brain activity decorrelates over development (*44, 45*). In line with these results, in our model, interconnected inhibitory neurons repel each other and their tuning curves decorrelate (Fig. 2*D*).

**Figure 2:**
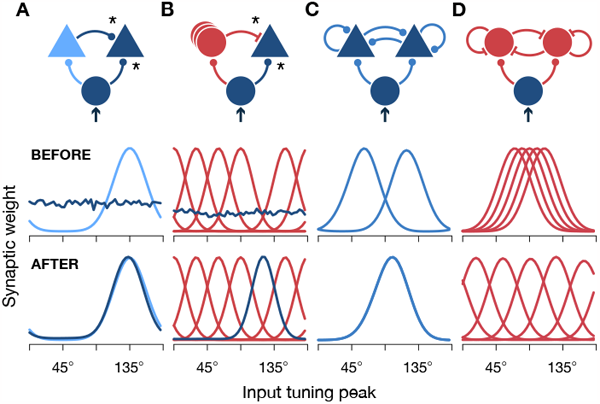
Feedforward tunings are affected by lateral input in microcircuit motifs. (*A*) In addition to feedforward input from a population of orientation tuned excitatory cells (blue circle), a neuron receives lateral input from an excitatory neuron with fixed feedforward tuning (light blue). * indicates plastic synapses. Feedforward tuning curves of the two neurons are shown before (center row) and after (bottom row) training. (*B*) Same as in *A*, for lateral input from multiple inhibitory neurons with fixed feedforward tuning. (*C*) Same as in *A*, for two recurrently connected excitatory neurons with all feedforward and recurrent synapses plastic. (*D*) Same as in *C*, for inhibitory neurons. All synapses plastic.

### Tuning curve decorrelation in fully plastic recurrent E-I networks

Recent experimental studies have suggested that inhibitory neurons drive decorrelation of neural activities (*46, 47*). Hence, we asked whether the interaction between excitatory and inhibitory neurons can serve to decorrelate not only inhibitory but also excitatory neural activities. To address this question we explored the consequences of synapse-type-specific competitive Hebbian learning in a network of recurrently connected excitatory and inhibitory neurons. We presented different oriented gratings in random order to a network where all feedforward and recurrent synapses are plastic (Fig. 3*A*, top). We observed a sharp increase in response selectivity (Fig. 3*A*, bottom) that is reflected in the reconfiguration of feedforward synaptic weights (cf. SM Movies M1 & M2): Shortly after random initialization (Fig. 3*B*, top), excitatory neurons predominantly connect to a subset of input neurons with similar stimulus selectivities (Fig. 3*B*, center left). We quantified the uniformity of the distribution of feedforward tuning curves during training (Fig. 3*C*, see Methods) and found that inhibitory neurons maintained a much wider coverage of the input stimulus space than the excitatory population (cf. Fig. 3*B*, center, *t*_1_). Eventually, tuning curves of excitatory as well as inhibitory neurons decorrelate and cover the whole stimulus space with minimal overlap (Fig. 3*B*, bottom), in sharp contrast to circuits without inhibition, where tuning curves become clustered (cf. Fig. 2*C*). After training, neurons are organized in an assembly-like structure. Neurons that are similarly tuned became more strongly connected (Fig. 3*D & E*), as is observed experimentally (*48*–*58*). We found that inhibitory neurons become as selective for stimulus orientations as excitatory neurons (*34*–*37*) (Fig. 3*F*), while excitatory input is balanced by similarly tuned inhibitory input (Fig. 3*G*) from multiple overlapping inhibitory neurons (Fig. 3*H*), in agreement with experimental results (*12, 59*– *64*); but see (*65*–*70*).

**Figure 3:**
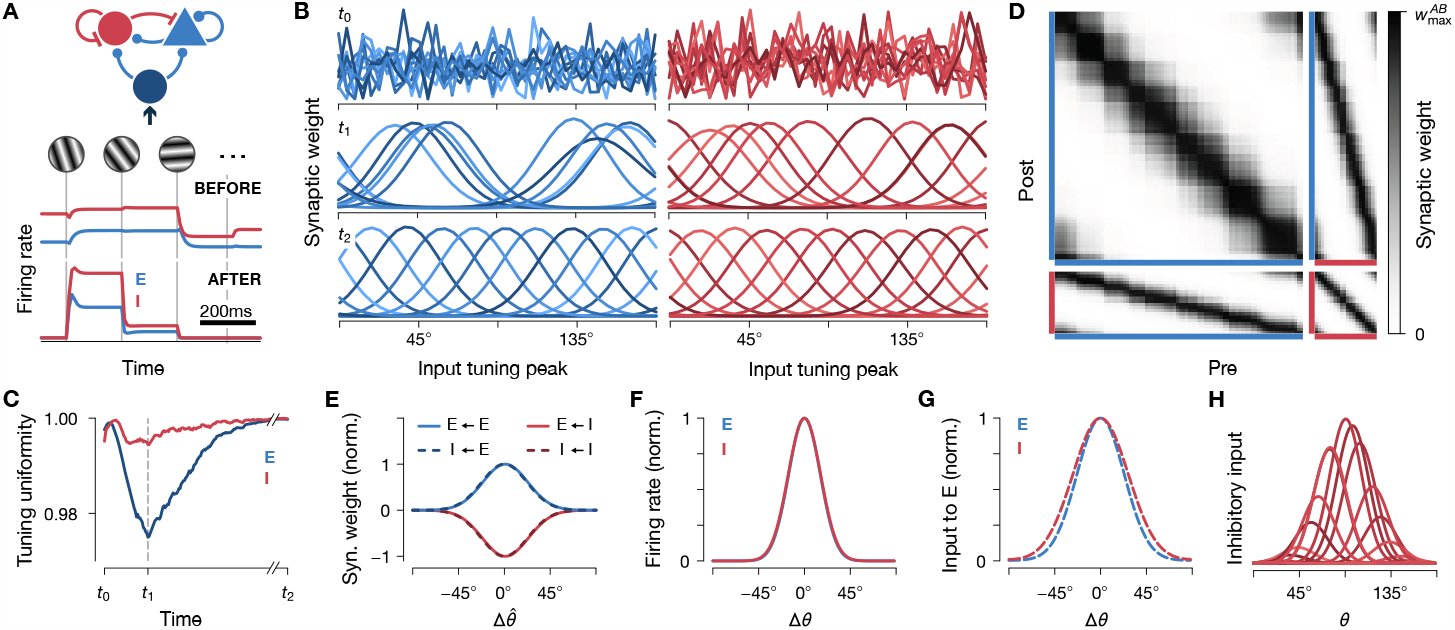
Tuning curve decorrelation in plastic recurrent networks. (*A*) Top: A population of recurrently connected excitatory and inhibitory neurons receives input from a set of input neurons that are tuned to different stimulus orientations (cf. Fig. 1*B*, bottom). Every 200ms a different orientation is presented to the network (vertical gray lines). At the same time, all synapses exhibit plasticity according to a synapse-type-specific Hebbian rule (see Methods). Bottom: typical firing rate activity of one excitatory (blue) and one inhibitory (red) neuron before and after training. (*B*) Feedforward tuning curves of *N*_*E*_ = 10 excitatory neurons before (*t*_0_, top), during (*t*_1_, center), and after (*t*_2_, bottom) training. Synaptic weights were initialized randomly. Different color shades indicate weights of different postsynaptic neurons. Compare SM Movies M1 & M2. (*C*) Feedforward population tuning uniformity (see Methods) of excitatory and inhibitory neurons in *B*. Time points *t*_0_, *t*_1_, *t*_2_ correspond to time points in *B*. (*D*) Connectivity matrices after training a network of *N*_*E*_ = 80 excitatory (blue) and *N*_*I*_ = 20 inhibitory (red) neurons. Neurons are sorted according to their preferred orientation 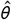, as measured by their peak response to different oriented gratings. 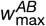 is the largest synaptic weight between population *A* and *B*; *A, B* ∈ {*E, I*}. (*E*) Normalized (norm.) recurrent weight strengths as a function of the difference between preferred orientations of the pre- and postsynaptic neurons, 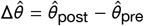, averaged over all neuron pairs. Input weights to excitatory (solid) and inhibitory (dashed) neurons overlap. (*F*) Average firing rate response of inhibitory and excitatory neurons to a stimulus orientation *θ*, relative to their preferred orientation, 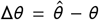, averaged over all neurons. Curves for excitatory (blue) and inhibitory (red) neurons overlap. (*G*) Same as in *F*, but for average excitatory and inhibitory inputs to excitatory neurons. (*H*) Inhibitory input to an excitatory neuron with preferred orientation close to 90°. Each curve corresponds to the input from one presynaptic inhibitory neuron for stimuli of different orientations *θ*.

In summary, we find that synapse-type-specific competitive Hebbian learning in fully plastic recurrent networks is sufficient to decorrelate neural activities and leads to preferential connectivity between similarly tuned neurons, as observed in cortical circuits.

### Inhibitory neurons balance excitatory attraction and enable decorrelation

To uncover how recurrent inhibition can prevent all neurons from becoming selective for a single input mode, we investigated the fundamental principles of synapse-type-specific competitive Hebbian learning in recurrent networks analytically (SM Sec. 5). In the simplified case of linear activation functions, input modes are eigenvectors of the input covariance matrix (cf. Eq.3). Since these eigenvectors are orthogonal by definition (Fig. 4*A*), the activities of neurons that are tuned to different eigenvectors are uncorrelated, and their reciprocal connections decay to zero under Hebbian plasticity (Fig. 4*B*). Then, neurons that are tuned to the same input mode form recurrent ‘eigencircuits’ that are otherwise separated from the rest of the network (SM Sec. 4). We characterize a mode’s effective attraction as a number such that, if a mode has a higher attraction than a competing mode, then neurons responding to the mode with lower attraction are unstable and shift their tuning towards the mode with higher attraction (see SM for details). Just like single, laterally projecting neurons (SM Sec. 3), eigencircuits also modify the effective attraction of their input mode (Fig. 4*C*). The decomposition of the network into eigencircuits allows to write the effective attraction *λ* of each input mode as the sum of a feedforward component *λ* and the variances of the neurons that reside in the respective eigencircuit (cf. SM Sec. 4.1 & 4.2):

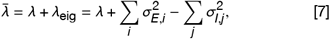

where we defined the contribution of recurrently projecting neurons to the effective attraction of an input mode as the eigencircuit attraction, *λ*_eig_. Note that, in general, variances 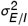 depend on the total synaptic weights, and the number of excitatory and inhibitory neurons in the eigencircuit (SM Sec. 4.2). This reveals that the attractive and repulsive effects of excitatory and inhibitory neurons can balance each other. In a simplified example, we assumed that all input modes have equal feedforward attraction, equal to *λ*, while each excitatory neuron contributes plus one and each inhibitory neuron minus one to the effective attraction (Fig. 4*D*, top). Then the eigencircuit attractions becomes *λ*_eig_ = *n*_*E*_ − *n*_*I*_ (Fig. 4*D*, bottom, solid line). In this configuration, the network is unstable: All neurons are attracted towards the input mode with the highest effective attraction (EC3), which suggests that all tuning curves will eventually collapse onto the same input mode. However, when all neurons become selective to the most attractive input mode, that mode would become repulsive (Fig. 4*D*, bottom, dashed gray line), as each increase in attraction due to an additional excitatory neuron is balanced by a decrease in attraction due to two additional inhibitory neurons. Consequently, the resulting eigencircuit is unstable and neurons are repelled towards non-repulsive, unoccupied input modes; distributed across the stimulus space.

**Figure 4:**
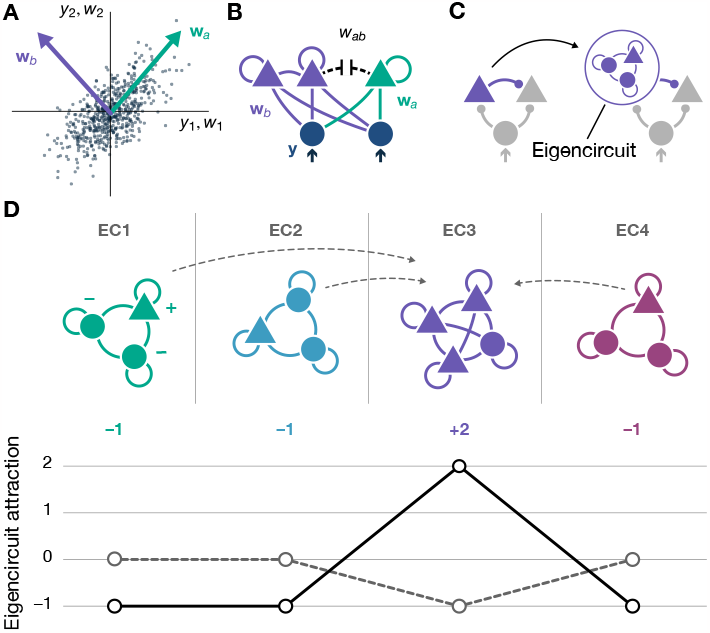
Illustration of eigencircuit decomposition and attraction. (*A*) Feedforward synaptic weight vectors **w**_*a*_, **w**_*b*_ of two neurons that are tuned to two different principal components (top, purple and green) of the input data. Each dark blue dot represents one presynaptic firing pattern (cf. Fig. 1*G*). (*B*) Synaptic weights *w*_*ab*_ between neurons that are tuned to different eigenvectors decay to zero, while neurons tuned to the same eigenvector form recurrently connected eigencircuits (purple). (*C*)As single, laterally projecting neurons shape the effective attraction of their input mode (left; cf. Fig.2), eigencircuits also increase or decrease the effective attraction of their respective eigenvector direction (right). (*D*) A recurrent network of excitatory (triangles) and inhibitory (circles) neurons that are distributed across four decoupled eigencircuits (EC, top). Each excitatory neuron contributes plus one (+), each inhibitory neuron minus one (-) to the eigencircuit attraction, *λ*_eig_ (solid line, bottom). Due to synaptic plasticity, neurons are pulled towards the most attractive eigencircuit, EC3 (gray dashed arrows, top). After all neurons integrate into the same eigencircuit (EC3), its attraction becomes negative, while the now unoccupied eigencircuits (EC1, EC2, EC4) are neutral (dashed line, bottom).

While this example conveys the core principle of how recurrently connected neurons adjust their tunings, the actual dynamics of synaptic weights are more complex (SM Sec. 5). In particular, neurons do not switch their tuning between input modes in discrete steps but shift their tuning gradually. Due to the recurrent nature of the circuit, even small tuning shifts affect the attractions of the respective eigencircuits (cf. SM Sec. 5.2.3). In our simulations, we therefore never observe a full collapse of all tuning curves onto the same input mode before neurons distribute across the stimulus space. Instead, neurons rapidly develop tuned feedforward receptive fields that gradually shift to maximise tuning uniformity, with little to no oscillatory dynamics (Fig. 3*B & C* and SM Movies M1 & M2).

In the simplified case of linear activation functions, we derive the following condition that prevents the collapse of all tuning curves onto a single input mode:

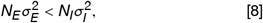

where 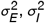 are the average of the variances of the excitatory and inhibitory firing rates, and *N*_*E*_, *N*_*I*_ are the total number of neurons in the network (cf. SM Sec. 5.2.4). These results show that recruiting recurrent inhibition can prevent tuning curve collapse and enables decorrelation, where a lower number of inhibitory neurons can be compensated by an increase in neural activation.

### Plastic recurrent E-I networks perform response normalization and exhibit winner-takes-all dynamics

Our results thus far reveal how synapse-type-specific competitive Hebbian learning can explain the development of structured recurrent connectivity. We next asked whether synapse-type-specific competitive Hebbian learning can also explain the emergence of non-linear network computations. For example, the firing rate response of neurons in the visual cortex to multiple overlayed oriented gratings is normalized in a non-linear fashion (*71, 72*). While this form of normalization is mostly of thalamic origin (*73*–*75*), there is most likely also a cortical component(*72, 76*). A recently introduced E-I network model with static, hand-crafted connectivity can explain these modulations (*18, 77*). We explored if the recurrent connectivity can instead be learned from a network’s input stimulus statistics. We consider a circuit with fixed feedforward tuning and plastic recurrent connectivity (Fig. 5*A*). After training the network with single oriented grating stimuli (Fig. 5*A*, bottom), we found that neural responses to a cross-oriented mask grating that is presented in addition to a regular test grating are normalized, i.e., the response to the combined stimulus is weaker than the sum of the responses to the individual gratings (Fig. 5*B*, left). When the contrast of the mask grating is lower than the test grating’s, the network responds in a winner-takes-all fashion: The higher-contrast test grating dominates activities while the lower-contrast mask grating is suppressed (Fig. 5*B*, right). As observed experimentally (*71, 72, 78*), we found that this orientation-specific response normalization is divisive and shifts the log-scale contrast-response function to the right (Fig. 5*C*).

**Figure 5:**
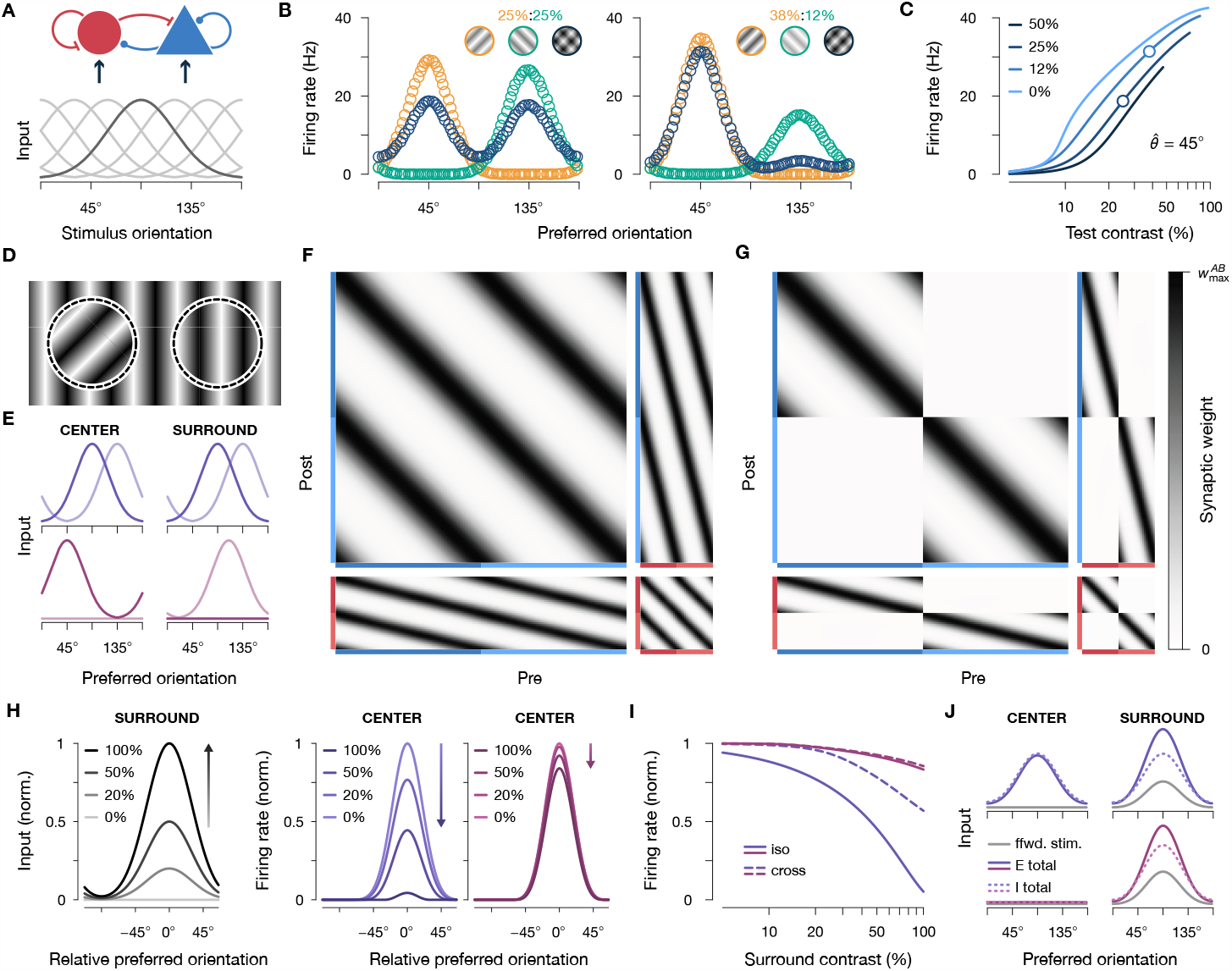
Cross-orientation and surround suppression in trained neural networks. (*A*) A plastic recurrent network of excitatory and inhibitory neurons (top) receives input according to fixed feedforward tuning curves (bottom). Input amplitudes were modulated with stimulus contrast. Tuning curve of neurons with preferred orientation of 90° highlighted in dark gray. (*B*) Response of 80 excitatory neurons to a test grating (orange, 45°) and a mask grating (green, 135°) of different contrast levels (insets, grating contrasts increased for better visibility). Gratings are presented separately (orange & green) or together (dark blue). Each open circle corresponds to the response of one excitatory neuron. (*C*) Contrast response curve of a single excitatory neuron with preferred orientation 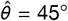 to the test and mask gratings in *B*. Different mask contrasts are indicated by different color shades. The bottom/top circles correspond to the left/right contrast level configurations in *B*. (*D*) Center (left) and surround region (right) with different oriented stimuli. (*E*) Example stimuli during training with different stimulus statistics. Top: Neurons tuned to the same orientation, but different regions (center region, left; or surround region, right) receive identical input; two example stimuli are shown in solid and transparent purple, respectively. Bottom: Neurons tuned to the center and surround regions are stimulated separately; two example stimuli are shown in solid and transparent red, respectively. Either the surround or the center regions are stimulated, while the other region receives zero input. (*F*) Recurrent connectivity matrix between excitatory (blue) and inhibitory (red) neurons (cf. Fig. 3*D*) after training the network with correlated center and surround stimuli (corresponds to purple color in *E*, top). Neurons are sorted according to their feedforward orientation tuning. Color shades indicate tuning to the center (dark) or surround (light) region. (*G*) Same as in *F*, but for a network trained with single gratings that were presented either in the center or the surround region (corresponds to red color in *E*, bottom). (*H*) Suppression of excitatory population activity in response to increasing surround stimulation for two networks trained under different stimulus statistics. Left: neurons tuned to the center region are stimulated by an oriented grating of constant, 100% contrast (not shown) while neurons tuned to the surround region are stimulated with an oriented grating of increasing contrast (shades; compare insets). Identical stimulation protocol for both training statistics. Center and right: the activity of excitatory neurons that are tuned to the center region is suppressed with increasing surround contrast. The magnitude of suppression depends on the stimulus statistics during training (purple vs. red, colors as in *E*). (*I*) Response of one excitatory neuron to center and surround stimulation after training. A center stimulus of preferred orientation was presented at constant contrast while the contrast of a cross- (dashed) or iso-oriented (solid) surround stimulus changed. Colors indicate different stimulus statistics during training (as in *E*). (*J*) Total excitatory (solid) and inhibitory (dotted) input to excitatory neurons during stimulation of only the surround region with an oriented grating of 90°. Excitatory input due to feedforward stimulation (ffwd. stim.) is shown in light gray. Colors (top vs. bottom) indicate different input statistics during training (as in *E*).

### Sensory input statistics shape computational functions of recurrent circuits

We next investigated how the stimulus statistics during training affect receptive field properties. We considered a plastic network where two neural populations receive tuned input from either a center or a surround region of the visual field (Fig. 5*D*). During training, we presented either the same oriented grating in both regions (Fig. 5*E*, top, purple), or a single grating in just one region (Fig. 5*E*, bottom, red), at 50% contrast (cf. Table 1). These stimulus statistics heavily influenced the recurrent connectivity structure in the network. When identical oriented stimuli are presented to the center and surround regions during training, neurons with similar orientation tuning become most strongly connected, independent from which region the neurons receive their feedforward input (Fig. 5*F*). However, when the center and surround regions are stimulated separately during training, neurons only connect to similarly tuned neurons within the same region and cross-region connectivity decays to zero (Fig. 5*G*). These differences in the recurrent connectivity structure are also reflected in the networks’ response properties. We found that after training, the response of center-tuned neurons exhibits orientation-specific surround suppression, reflecting the stimulus statistics during training. When the center and the surround regions are stimulated separately during training, iso- or cross-oriented stimuli in the surround both elicit minimal suppression of the center-tuned population’s response to a center stimulus (Fig. 5*H & I*, red). In contrast, in the case of correlated stimulation of the center and surround regions during training, the response of the center population is markedly suppressed when an additional surround stimulus is presented (Fig. 5*H & I*, purple). Importantly, suppression is stronger for iso-compared to cross-orientations (Fig. 5*I*, solid and dashed lines), as has been reported experimentally (*79*–*82*). We further investigated the lateral interactions between neurons tuned to the center and surround regions by presenting an oriented stimulus only in the surround region, while observing the total excitatory and inhibitory inputs to excitatory neurons (Fig. 5*J*). We found that the total excitatory input to stimulated excitatory neurons in the surround was larger than the total inhibitory input (Fig. 5*J*, right column). When center and surround neurons were stimulated together during training, both center and surround received similar, balanced E and I recurrent input, but the surround cells also received feedforward excitation, yielding more total excitation (Fig. 5*J*, top, purple). When center neurons were not stimulated with the surround neurons during training, they received no input from a surround-only stimulus (Fig. 5*J*, bottom, red). In the case of correlated stimulation of the center and surround regions during training, this lateral input was orientation-specific. Center neurons tuned to the same orientations as stimulated neurons in the surround received stronger input than center neurons tuned to different orientations (Fig. 5*J*, top left), reflecting the input stimulus statistics during training (Fig. 5*E*) and the resulting recurrent connectivity (Fig. 5*F*). A similar balance of excitatory and inhibitory lateral inputs has previously been observed in barrel cortex (*83*). Together, this demonstrates that synapse-type-specific competitive Hebbian learning produces extra-classical receptive fields that modulate feedforward responses via recurrent interactions that reflect the input statistics during training.

**Table 1:** Simulation parameters. Different parameters for different panels are separated by commas. Dashes indicate that parameters were not used in the simulation. For Fig. 5, weight norms of static synaptic weights are indicated by ‘‡’. For Fig. 2, parameters of static, laterally projecting neurons are given in brackets (comma separated for panels *A, B*).

| Figure | 1E-F, H-I | 2A, B, C, D (A, B) | 3A-C, D-H | 5A-C, D-J |
| --- | --- | --- | --- | --- |
| $N_E$ | 1 | 2, 1, 2, 0 | 10, 80 | 80, 80 × 2 |
| $N_I$ | 10 | 0, 5, 0, 5 | 10, 20 | 20, 20 × 2 |
| $N_F$ | 10 | 50 | 40, 80 | 80, 80 × 2 |
| $a$ | 1 | 0.04 (0.2, 0.08) | 0.04 | 0.04 |
| $b$ | 0.25 | 0 | 0 | 0 |
| $n$ | 2 | 2 | 2 | 2 |
| $\mu_W$ | 0.1 | 0.1 | 0.2 | 0.2 |
| $\sigma_W$ | 0.05 | 0.01 | 0.1 | 0.1 |
| $W_{EE}$ | 10 | 1, 1, 4, - | 2, 0.6 | 3.51 |
| $W_{IE}$ | - | - | 2, 0.85 | 3.35 |
| $W_{EI}$ | 5, - | -, 0.5, -, - | 0.8, 0.3 | 1.84 |
| $W_{II}$ | - | -, -, -, 0.5 | 0.5, 0.35 | 1.44 |
| $W_{EF}$ | - | (0.9, -) | - | 1.4<sup>‡</sup> |
| $W_{IF}$ | - | (-, 1) | - | 1.4<sup>‡</sup> |
| $c$ | 1 | 1 | 1 | 0.5 |
| $A_F$ | 1 | 1 | 35, 140 | 80 |
| $\sigma_F$ | 20° | 12° | 12° | 30°/√2 |
| $\sigma_\theta$ | - | -, -, 22°, 22° (22°, 16°) | - | 30°/√2 |
| $\Delta t$ | 200ms | 20ms | 10ms | 10ms |
| $\tau_E$ | 200ms | 40ms | 20ms | 25ms |
| $\tau_I$ | - | 28ms | 17ms | 12.5ms |
| $\epsilon_{EE}$ | - | $0.4 \times 10^{-8} \text{ms}^{-1}$ | $2 \times 10^{-9} \text{ms}^{-1}$, $1.0 \times 10^{-10} \text{ms}^{-1}$ | $1.0 \times 10^{-9} \text{ms}^{-1}$ |
| $\epsilon_{IE}$ | - | $0.8 \times 10^{-8} \text{ms}^{-1}$ | $3 \times 10^{-9} \text{ms}^{-1}$, $1.5 \times 10^{-10} \text{ms}^{-1}$ | $1.5 \times 10^{-9} \text{ms}^{-1}$ |
| $\epsilon_{EI}$ | $2 \times 10^{-4} \text{ms}^{-1}$, $4 \times 10^{-4} \text{ms}^{-1}$ | $0.6 \times 10^{-8} \text{ms}^{-1}$ | $4 \times 10^{-9} \text{ms}^{-1}$, $2.0 \times 10^{-10} \text{ms}^{-1}$ | $2.0 \times 10^{-9} \text{ms}^{-1}$ |
| $\epsilon_{II}$ | - | $1.0 \times 10^{-8} \text{ms}^{-1}$ | $5 \times 10^{-9} \text{ms}^{-1}$, $2.5 \times 10^{-10} \text{ms}^{-1}$ | $2.5 \times 10^{-9} \text{ms}^{-1}$ |
| $\epsilon_{EF}$ | $1 \times 10^{-4} \text{ms}^{-1}$, $2 \times 10^{-4} \text{ms}^{-1}$ | $\epsilon_{EE}$ | $\epsilon_{EE}$ | - |
| $\epsilon_{IF}$ | - | $\epsilon_{IE}$ | $\epsilon_{IE}$ | - |
| $r_0$ | -, 0.25 | - | - | - |

## Discussion

Our results suggest that synapse-type-specific competitive Hebbian learning is the key mechanism that enables the formation of functional recurrent networks. Rather than hand-tuning connectivity to selectively explain experimental data, our circuits emerge from a single unsupervised, biologically plausible learning paradigm that acts simultaneously at all synapses. In a single framework, our networks readily explain multiple experimental observations, including the development of stimulus selectivity, excitation-inhibition balance, decorrelated neural activity, assembly structures, response normalization, and orientation-specific surround suppression. These results demonstrate how the connectivity of inhibition-balanced networks is shaped by their input statistics and explain the experience-dependent formation of extra-classical receptive fields (*84*–*88*). Unlike previous models (*89*–*94*), our networks are composed of excitatory and inhibitory neurons with fully plastic recurrent connectivity.

Early theoretical work on inhibitory plasticity assumed that synapses evolve to maintain the mean firing rate of postsynaptic excitatory neurons (*23*). When excitatory input is static, this leads to neural tunings where inhibition and excitation are balanced. However, when excitatory are simultaneously plastic according to a simple Hebbian rule, the circuit is unstable and can not explain the joint development of feedforward stimulus tuning and inhibitory balance (*24*) (SM Sec. 1.2.3). The system can be stabilized when the Hebbian growth of excitatory synapses is controlled by a BCM-like plasticity threshold. This introduces fierce competition between different input streams in the form of subtractive weight normalization, which leads to winner-takes-all dynamics among synapses that do not allow for the development of extended receptive fields (*24, 31, 95*). Later models have proposed more intricate plasticity rules, some of which consider, e.g., voltages or currents, in addition to pre- and postsynaptic action potentials (*25, 28, 96*–*102*), as summarized in several recent reviews (*14, 103*–*106*). In recent years, there has also been a resurgence of interest in normative approaches (*28, 29, 107*). In these approaches, it is postulated that synaptic plasticity rules act to optimize an objective function that describes a desirable network property. Motivated by the notorious instability of recurrent networks, one obvious objective is stability, e.g., in the form of firing rate homeostasis. Following early theoretical work that suggested such a homeostatic role for synaptic plasticity of inhibitory synapses onto excitatory neurons(*23*), two recent studies propose a similar role for the plasticity of other recurrent synapse types (*28, 29*). Indeed, such plasticity rules allow the formation of inhibition balanced receptive fields (*28*), and stabilize network activity, even when faced with strong recurrent connections (*29*). However, none of these rules have been applied in fully plastic recurrent networks with structured feedforward input. Even in complex models that use many different forms of plasticity, some synapse types are kept static after initialization, to maintain stable network activity (*23, 26, 27, 101*). While such networks still show many interesting dynamics, they lack the rich computational functions of circuits with structured connectivity between all neuron types (*18, 77*). In contrast, our learning rule is minimalistic and only relies on general Hebbian synaptic growth that is stabilized by competitive interactions. Importantly, our theory does not depend on a specific biophysical implementation of the Hebbian plasticity paradigm. We only require that synapses follow the basic Hebbian principle of synaptic strengthening following concurrent pre- and post-synaptic activity. In the past, competitive Hebbian learning has been investigated theoretically for excitatory synaptic inputs to single neurons (*30, 31, 39, 108, 109*), but not for inhibitory inputs or in recurrent networks. Our analysis demonstrates that competitive Hebbian plasticity is a suitable learning mechanism for networks of recurrently connected excitatory and inhibitory neurons, while being analytically tractable and biologically plausible.

Competitive interactions between synapses have been observed in many different preparations and have been attributed to various mechanisms (*110*–*121*). While previous work has focused on competitive interactions between excitatory synapses, our results support the notion that similar competitive processes are also active at inhibitory synapses (*32, 122*). The local competition for a limited supply of synaptic building blocks is a biologically plausible normalization mechanism (*33, 115, 120, 123, 124*). Many synaptic proteins are specific to inhibitory or excitatory synapses and reside in one synapse-type, but not the other (*125, 126*). Therefore, in this work, we assume a synapse-type-specific competition for different synaptic resource pools and implement separate normalization constants for inhibitory and excitatory synapses. On a finer scale, synapses of different excitatory and inhibitory neuron subtypes also differ in their protein composition (*126*–*129*). In principle, this allows for the precise regulation of different input pathways via the adjustment of subtype-specific resource pools (*130*–*136*). Furthermore, axons of different neuron sub-types target spatially separated regions on the dendritic tree, allowing for pathway-specific local competition. For example, somatostatin-positive cortical Martinotti cells target the apical dendritic tree of pyramidal cells, while parvalbumin-positive basket cells form synapses closer to the soma (*1*), which suggests that afferents of these cell types compete for separate resources pools. We anticipate such subtype-specific mechanisms to be crucial for the functional development of any network with multiple neuron subtypes (*137, 138*).

In the brain, total synaptic strengths are dynamic and homeostatically regulated on a timescale of hours to days (*139*–*142*). In addition to maintaining average firing rates in response to network-scale perturbations, a prominent framework puts forward homeostatic scaling of synaptic strengths as a stabilizing mechanism of Hebbian growth (*143*). However, theoretical models suggest that homeostatic scaling is too slow to balance rapid synaptic plasticity (*144*). In our networks, Hebbian growth is instead thought to be stabilized by the competition for a limited pool of synapse-type-specific resources, while total synaptic strengths remain fixed. This competition is fast due to rapid interactions on a molecular level (*33, 120*). Compared to Hebbian growth, infinitely fast, as a synapse can only grow at the expense of another. Therefore, we suggest that homeostatic scaling of total synaptic strengths is not required for immediate network stability but instead controls the operating regime of the network (*16, 77, 145*).

Our results demonstrate how multi-synaptic, inhibitory interactions can decorrelate excitatory neurons. In contrast, inhibitory neurons can inhibit each other mono-synaptically and do not require additional recurrent interactions for decorrelation. Accordingly, we observe that during training, inhibitory neurons are more decorrelated compared to excitatory neurons (Fig. 3*C*). These insights complement recent experimental results that suggest an instrumental role of inhibition in the decorrelation of excitatory networks in mouse prefrontal cortex during early development (*47*). Recent experimental studies in ferret visual cortex report conflicting evidence — either supporting (*46*) or contradicting (*146*) aligned developmental trajectories of excitatory and inhibitory populations. In our simulations, we observe similar developmental trajectories for excitatory and inhibitory populations. However, we focused on synaptic plasticity and did not consider other processes, like critical periods (*147, 148*), that are known to shape circuit development.

Cortical computations rely on strong recurrent synaptic weights that result in neural activities that can deviate significantly from the input stimulus pattern (*15, 16, 18*) (cf. Fig. 5*B*, left, combined grating response). Such a decoupling of network activity from feedforward input due to recurrent interactions can lead to neural tunings that do not reflect the input stimulus statistics (cf. SM Sec. 3). In our theory (SM Sec. 4), we assume that neurons are tuned to feedforward modes and thereby implicitly assume that network activity is dominated by feedforward input. In our numerical simulations of fully plastic recurrent networks, we find that for intermediate levels of recurrence (cf. Table 1, Fig. 1, 2 & 3), the network’s activities are indeed dominated by feedforward inputs. In case of strong recurrence (Fig. 5), we ensure feedforward dominance by presenting single oriented gratings that match the fixed feedforward tunings of neurons (cf. Fig. 5). Such gratings elicit a Gaussian-shaped response that is sharpened due to the recurrent connectivity, but maintains the general correlation structure compared to purely feedforward-driven networks (compare tuning widths in Fig. 5*A*, bottom, and *B*, single grating response). Biological cortical networks are strongly recurrently connected (*149*–*153*). However, neural activity and the induction and polarity of synaptic plasticity are regulated by neuromodulators (*154*–*158*), which may control the destabilizing effect of strong recurrent connectivity. In addition, different synapse types do not develop simultaneously but progress through different developmental stages (*137, 159, 160*). For example, the development of recurrent excitatory connections is delayed compared to that of feedforward synapses (*131, 161*). Taking these factors into account will be essential for future models of developing recurrent circuits.

In our networks, structured feedforward input is crucial for the development of orientation selective receptive fields. However, already at the time before eye opening cortical neurons exhibit substantial selectivity for stimulus orientation, without having been exposed to the statistical regularities of visual inputs (*162*–*164*). One hypothesis is that, instead, spontaneous activity in the retina provides the statistical structure required for the initial development of orientation selectivity (*165*–*167*). In our model, circuit formation depends only on the statistical regularities between input streams and is agnostic with respect to their origin. Therefore, we expect our approach to extend beyond sensory cortices and to provide a fundamental framework for plasticity in recurrent neural networks.

## Materials and Methods

### Computational model

We consider networks of rate coding excitatory (*E*) and inhibitory (*I*) neurons that receive input from themselves and a population of feedforward input neurons (*F*). Membrane potential vectors **u** evolve according to

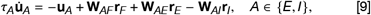

where *τ*_*A*_ is the activity timescale. **W**_*AB*_ are matrices that hold synaptic weights between the presynaptic population *B* and the postsynaptic population *A* with *B* ∈ {*E, I, F* }. All differential equations were numerically integrated using the Euler method in timesteps of Δ*t*. Entries of weight matrices were drawn from a normal distribution with mean *μ*_*W*_ equal to two times the standard deviation *σ*_*W*_, which yields mainly positive entries. Negative entries were set to their absolute value. Before the start of the simulation, excitatory and inhibitory weights were normalized as described below. Unless stated otherwise, prior to normalization, all *recurrent* excitatory and inhibitory weights were set to zero, i.e., initialized networks were purely feedforward. Firing rate vectors **r**_*A*_ are given as a function *f* (**u**_*A*_) of the membrane potential **u**_*A*_:

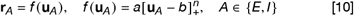

with [ · ]_+_ = max(0, ·) and scalar constants *a, b*, and *n*.

### Plasticity and normalization

Plastic weights evolve according to a Hebbian plasticity rule

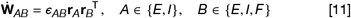

where ε_*AB*_ is a scalar learning rate, and^T^ indicates the transpose. After each plasticity step, synaptic weights are normalized such that the total excitatory and inhibitory postsynaptic weights are maintained:

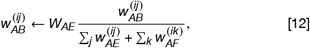

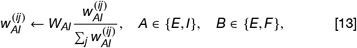

where *W*_*AE*_, *W*_*AI*_ are the total excitatory and inhibitory synaptic weight norms. Weights are updated and normalized in every integration timestep Δ*t*, in sync with the network dynamics.

In Fig. 1, we set the activity of the inhibitory input neurons equal to the activity of the excitatory input neurons, i.e., **r**_*I*_ = **r**_*F*_. For panels *H & I* of Fig. 1, inhibitory weights evolved according to the classic inhibitory plasticity rule (*23*) without normalization:

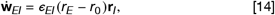

where *r*_0_ is a target firing rate.

### Input model

The activity of feedforward input neurons depends on the orientation *θ* and contrast *c* of an input grating:

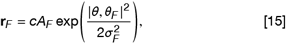

where the vector *θ*_*F*_ holds the preferred orientations of the input neurons that are evenly distributed between 0 and 180°, *σ*_*F*_ is the tuning width, *A*_*F*_ the maximum firing rate, and |·,· | is the angular distance, i.e., the shortest distance around a circle of circumference 180°. During training, single gratings, sampled from a uniform distribution between 0° and 180°, were presented to the network for 200ms, before the next stimulus was selected.

In Fig. 5 network stimulation is realized via static feedforward weights. Neuron were assigned a preferred orientation 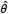, evenly distributed between 0° and 180°. Static feedforward weights were initialized as

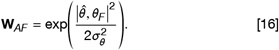

For Fig. 5, feedforward weights are normalized separately to *W*_*AF*_ before the start of the simulations (cf. Table 1). In this case, feedforward weights are fixed and are not taken into account when normalizing recurrent weights. Feedforward weights of static neurons in Fig. 2*A & B* are processed in the same fashion. For Fig. 5, parameters were selected to result in stimulation patterns as in Rubin *et al*. (*18*). Weight norms *W*_*AB*_ were also adapted from Rubin *et al*. (*18*). See Table 1 for an overview of used simulation parameters.

### Tuning curve uniformity measure

In Fig. 3*C*, we quantified the uniformity of the distribution of tuning curves during learning and defined:

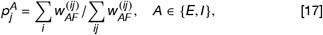

where 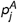is the normalized total synaptic output weight of input neuron *j* onto the excitatory (*E*) and inhibitory (*I*) neural population. Then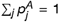, and we can define the tuning uniformity *U*_*A*_ as the normalized Shannon entropy 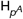.

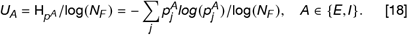

*U*_*A*_ is maximal, equal to one, if *p*^*A*^ is uniformly distributed, and minimal, equal to zero, if all synaptic weight is concentrated in a single input neuron. Such a concentration is highly unlikely. In our simulations, weight distributions are much closer to a uniform distribution, and the uniformity measure is close to one.

## Acknowledgements

We thank the members of the Computation in Neural Circuits group for helpful feedback throughout the project. We thank Dylan Festa for additional comments and for proofreading the manuscript. This work was supported by the Max Planck Society, the Engineering and Physical Sciences Research Council (DTP grant EP/T517847/1 to E.J.Y), and the European Research Council (StG 804824 to J.G.).).

## Author contributions

SE conceived research with input from JG. SE performed simulations, derived mathematical results, prepared figures, and wrote the Supplementary Material and the first draft of the manuscript. EJY derived mathematical results and contributed to the Supplementary Material. SE and JG wrote the manuscript.

## Supplementary material

Supplementary material includes: SM Text Sections 1 to 6, Figures S1 to S4, Movies M1 and M2. Python code to reproduce the key results of the study is publicly available on GitHub (*168*).

## Supplementary Material for

### 1 Linear competitive Hebbian learning finds principal components

Before considering inhibitory plasticity, we recapitulate how linear Hebbian learning finds the principal eigenvector of a neuron’s inputs. Although first described by Oja (*1*), we will mostly follow the derivation by Miller and MacKay (*2*) that we will later extend to inhibitory neurons.

#### 1.1 Hebbian plasticity without normalization is unstable

We consider a single neuron that receives input from a set of excitatory neurons (Fig. S1*A*). Its output firing rate *r* is a weighted sum of the firing rates of its pre-synaptic inputs **y**. One can conveniently write this as a dot product:

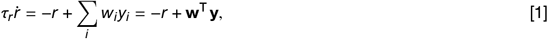

where **w** is a vector that holds the synaptic weights, and *τ*_*r*_ defines the timescale at which the activity changes. In the following, lowercase letters in bold indicate vectors, and uppercase letters in bold matrices. Following Hebb’s principle, synaptic weight changes depend on the pre- and post-synaptic firing rates. In vector notation:

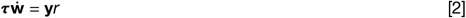

where the constant *τ* sets the timescale of plasticity. Assuming that synaptic weights change on a much slower timescale than firing rates, *τ*_*r*_ « *τ*, we make the simplifying assumption that *r* reaches its fixed point instantaneously, i.e., *τ*_*r*_ = 1 and *r* = **w**^⊤^**y**, and consider the same plasticity timescale for all synapses ***τ*** = 𝟙. Then, the average change of the synaptic weights can be expressed as a linear transformation of the original weight vector:

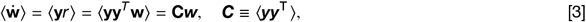

where ⟨ · ⟩is a temporal average and ***C*** is the covariance matrix of the synaptic inputs **y**, assuming inputs have zero mean, ⟨**y**⟩ = **0**. In the following, we only consider the average weight changes and omit the angled notation for convenience. To solve this differential equation, we express the weight change in the eigenvector basis of the covariance matrix **C**, which is symmetric and positive-semidefinite and, therefore, has a complete set of orthonormal eigenvectors with non-negative eigenvalues.

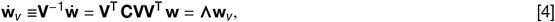

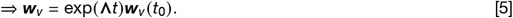

Here, **Λ** is the diagonal eigenvalue matrix, and each column of **V** holds mutually orthogonal eigenvectors, i.e., **VV**^⊤^ = 𝟙, and **V**^−1^ = **V**^⊤^. Each eigenvector component grows exponentially at a rate given by the respective eigen-value, which we identify with the *attraction* of the input component. We call eigenvector components with positive eigenvalue *attractive*, and the eigenvector component with the largest eigenvalue the *most attractive* input mode. We will later see that eigenvalues that describe the dynamics of input modes can become negative (Sec. 2). We will call such input modes with negative corresponding eigenvalue *repulsive*.

In summary, we find that unconstrained Hebbian plasticity results in the unlimited growth of synaptic weights and is therefore unstable. One way to constrain this unlimited growth is to modify the Hebbian learning rule such that the total synaptic weight is maintained.

**Figure S1:**
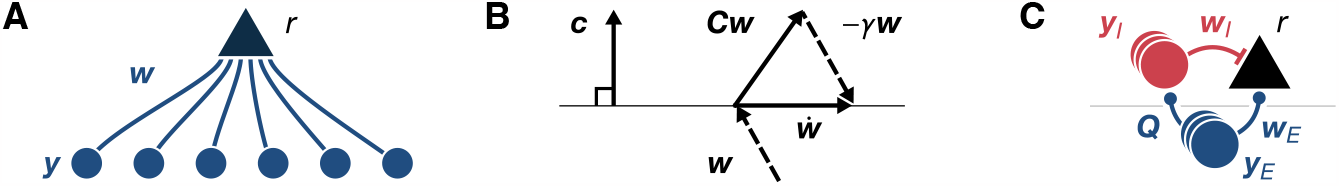
(*A*) Feedforward excitatory circuit. A post-synaptic neuron with output firing rate *r* receives synapses ***w*** from a set of excitatory neurons with firing rates *y*_*E*_. (*B*) The normalization operation constrains synaptic weight changes 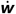 to a hyperplane that is perpendicular to the constraint vector ***c*** by subtracting a multiple *γ* of the weight vector ***w***. See text for details. Figure adapted from Miller and MacKay (*2*). (*C*) Feedforward inhibitory circuit. A post-synaptic neuron with output firing rate *r* receives excitatory synapses ***w***_*E*_ from a population of *N*_*E*_ excitatory neurons with firing rates ***y***_*E*_, and inhibitory synapses ***w***_*I*_ from a population of *N*_*I*_ inhibitory neurons with firing rates ***y***_*I*_. The gray horizontal line indicates the separation between two hypothetical brain regions or cortical layers.

#### 1.2 Weight constraints stabilize unlimited Hebbian growth

Hebbian plasticity and weight normalization can be considered as two discrete steps. First, growing weights according to the Hebbian rule. Second, normalizing to maintain the total synaptic weight. In this section, we will follow Miller and MacKay (*2*) and show how one can integrate these two discrete steps into one and derive the effective weight change 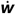. One can write the two steps as

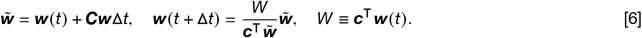

This update rule maintains the projection of ***w*** onto the constraint vector ***c*** by multiplicatively scaling the weight vector after the Hebbian learning step, i.e., 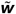. Alternatively, if we let *W* be a constant, the projection onto ***c*** would be constrained to be equal to that constant. In the following, we instead assume that the weights are already properly normalized and set the projection value as it was before the plasticity timestep, i.e., equal to *W* as defined above.

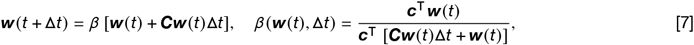

where *β* describes the multiplicative normalization that depends on the size of the timestep Δ*t* and the previous weight *w* (*t*). It is straightforward to check that the projection of the weight vector onto the constraint vector ***c*** does not change, i.e.,

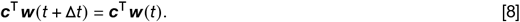

Then, the effective weight change 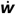 is given as

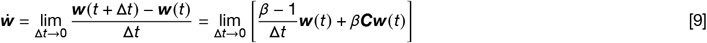

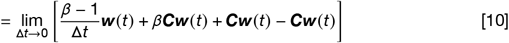

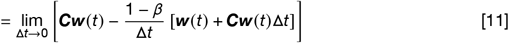

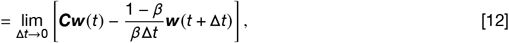

where, in the first and last steps, we used the definition of ***w***(*t* + Δ*t*) in Eq. 7. Next, we take the limit

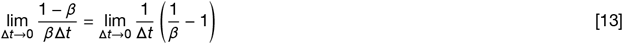

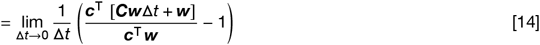

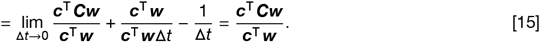

In summary, we get (cf. Fig. S1*B*):

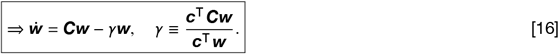

Here, *γ* is a scalar normalization factor that depends on the current weight ***w***.

An alternative way to derive 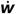 is to guess the shape of the multiplicative normalization term in Eq. 16 and require that the change along the constraint vector is zero^1^, i.e.,

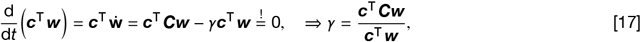

Note that for ***c*** being a constant vector of ones, the L1-norm of the weight vector is maintained. However, ***c*** does not have to be constant. For example, for ***c*** = ***w*** the L2-norm is maintained. Also, note that one can analogously derive effective plasticity rules when weights are constrained via subtractive normalization with the ansatz 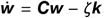 where ***k*** is a vector of ones (*2*).

##### 1.2.1 Fixed points

From Eq. 16 it is clear that all fixed points, for which 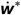 = 0, are multiples of eigenvectors ***v*** of ***C***. Explicitly, for a scalar constant *a* and ***w***^*^ = *a****v*** one gets:

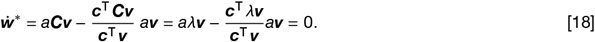

Note that this is independent of the choice of the constraint vector ***c***. We next consider the stability of these fixed points.

##### 1.2.2 Stability analysis

In the previous sections, we showed how multiplicative normalization constrains the norm of the weight vector and therefore prevents the otherwise unlimited growth of Hebbian plasticity. However, even when the total synaptic weight is constrained, synaptic weights might still be unstable and never settle into a fixed point, e.g., experiencing oscillatory dynamics and unstable fixed points. Following Miller and MacKay (*2*), we will now explore under what conditions fixed points are stable.

Formally, a fixed point in a linear system is stable when the largest eigenvalue of the Jacobian is negative, or marginally stable when it is equal to zero (*3*). The weight dynamics around a fixed point ***w***^*^ can be approximated with its Taylor expansion:

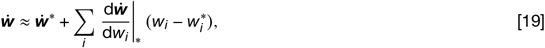

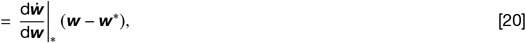

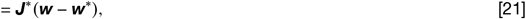

where 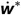 is zero, by definition, and ***J***^*^ is the Jacobian evaluated at the fixed point. The Jacobian is defined as

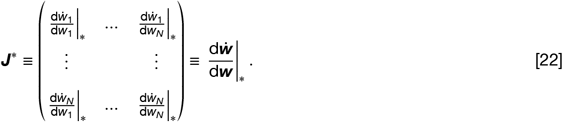

A fixed point is stable if small perturbations away from the fixed point, Δ***w*** = ***w*** − ***w***^*^, decay to zero, i.e.,

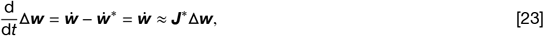

where we approximated 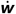 with its Taylor expansion (Eq. 19), since the perturbation is small, i.e., ***w*** is close to the fixed point. The result is a linear differential equation that one can solve as

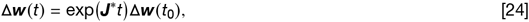

where all vector components decay to zero if all eigenvalues of ***J***^*^ are negative^1,2^. As we will see later, it is useful to rewrite the weight dynamics (Eq. 16) as

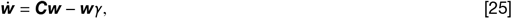

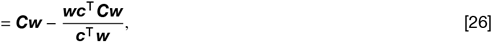

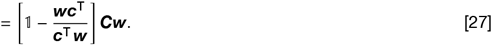

It follows^1^:

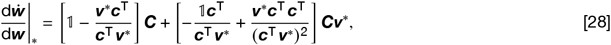

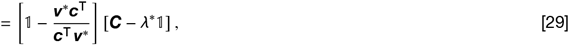

where ***w*** |_*_ = ***w***^*^ = *a****v***^*^ is the fixed point with ***v***^*^ being an eigenvector of ***C***. The scalar *a* is the length of the fixed point weight vector ***w***^*^ (which cancels) and *λ*^*^ is the eigenvalue to ***v***^*^. To find the eigenvalues of the Jacobian, *λ*_***J***_, we diagonalize ***J*** by switching to the eigenbasis of ***C***. When ***V*** is the matrix that holds the eigenvectors of ***C*** as columns one gets

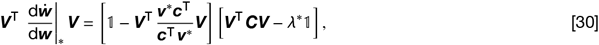

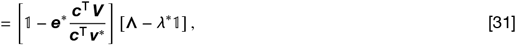

where **Λ** is a diagonal matrix that holds the eigenvalues of ***C***. Without loss of generality, we can assume that the first column of ***V*** is equal to ***v***^*^. Then ***e***^*^ = ***V***^⊤^***v***^*^ is a column vector of zeros, except for the first entry, which is equal to one. Then, the first bracket becomes an upper triangular matrix with ones on the diagonal, except for the first diagonal entry, which is zero. From this, it follows^2^ that the eigenvalues of the Jacobian are

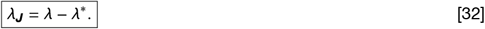

If *λ*^*^ is the largest eigenvalue, i.e., ***w***^*^ is a multiple of the principal eigenvector of ***C***, then all *λ*_***J***_ are negative or zero, and the fixed point is marginally stable. If there exists a *λ* > *λ*^*^, the corresponding *λ*_***J***_ is positive and the fixed point is unstable. Therefore, the eigenvector corresponding to the principal eigenvalue is the only (marginally) stable fixed point. In summary, linear Hebbian learning combined with multiplicative normalization becomes selective for the principal eigenvector of the input covariance matrix and thus performs principal component analysis (PCA). Next, we consider what happens when a neuron also receives inhibitory input.

##### 1.2.3 Classic Inhibitory plasticity prevents stimulus selectivity

Previous work suggested a homeostatic inhibitory synaptic plasticity rule (*4*) that enforced a post-synaptic target firing rate *r*_0_:

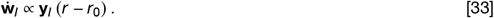

However, when combined with excitatory plasticity, this classic rule prevents the development of stimulus selectivity. For completeness, we briefly recapitulate this result presented in Clopath *et al*.(*5*): The authors find that classic inhibitory plasticity is required to act faster than excitatory plasticity to enable stable weight dynamics (*5*). For much faster inhibitory plasticity, the dynamics of excitatory and inhibitory weights decouples, and fixed points of the inhibitory weights 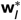 can be considered separately from the fixed points of excitatory weights. When excitatory and inhibitory inputs are equally stimulus selective, the fast dynamics of inhibitory weights ensures that the target firing rate is consistently met, i.e., the post-synaptic neuron always responds with the same firing rate *r*^*^.

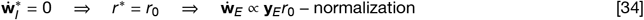

When all pre-synaptic neurons have similar average firing rates, ⟨*y*_*E*_⟩ _*i*_ ≈*y*_0_, and weights change on a slower timescale than activities as is the case biologically, the average excitatory synaptic weight change becomes

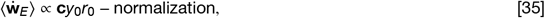

where **c** is a vector of ones. The average synaptic weight change is identical across synapses, which prevents the development of stimulus selectivity (Fig. 1*E & F*). Therefore, classic inhibitory plasticity that enforces a target firing rate cannot explain the joint development of stimulus selectivity and inhibitory balance. Instead, we propose that, as excitatory weights, also inhibitory weights are constrained via a competitive process that normalizes the total inhibitory input a neuron receives.

### 2 Synapse-type-specific normalization balances E-I receptive fields

Different from the normalization of excitatory weights, the normalization of inhibitory weights is not motivated by the requirement for stability. Inhibitory synaptic plasticity that depends on neural activity is self-limiting, since increasing inhibitory weights eventually prevent the neuron from firing, and thus prevent further plasticity. Instead, we motivate the normalization of inhibitory synaptic weights by the competition for a limited amount of synaptic building blocks that may also drive excitatory normalization (see Main text for details).

In the following, we generalize the approach outlined above for excitatory weight normalization to the case of simultaneous excitatory and inhibitory normalization. We consider a simplified circuit of a single post-synaptic neuron with firing rate *r* that receives lateral input from *N*_*I*_ inhibitory neurons, while all neurons receive feedforward input from a population of *N*_*E*_ excitatory neurons^1^. Then ***y***_***E***_ and ***y***_***I***_ are vectors that hold the firing rates of the excitatory and inhibitory populations. We now explore the self-organization of excitatory and inhibitory synaptic weights, ***w***_*E*_ and ***w***_*I*_, that project onto the single post-synaptic neuron, while input synapses ***Q*** that project onto inhibitory neurons remain fixed (Fig. S1*C*). Then, the rate dynamics reads

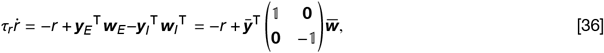

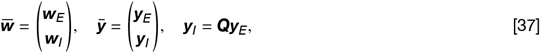

where 1 is the unit matrix with appropriate dimension, **0** are matrices of zeros and appropriate dimensionality, and we defined the modified weight and input vectors, 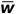 and 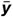. Similar to before, we assume fast activity dynamics, *τ*_*r*_ = 1, and write the Hebbian part of the time-averaged weight dynamics as

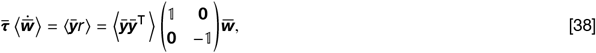

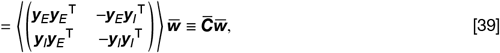

where we defined the modified covariance matrix 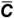. In general, we assume that all synapses of one type, excitatory or inhibitory, change equally fast (cf. Table 1). Then, the matrix 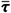 holds the timescales of excitatory plasticity, ***τ***_*E*_ = 𝟙, *τ*_*E*_, and inhibitory plasticity, ***τ***_*I*_ = 𝟙 *τ*_*I*_, as matrices on the diagonal, and is zero otherwise. In the following, we drop the bracket notation⟨ · ⟩ for better readability. As in the case of only excitatory input, we can implement multiplicative normalization by additional constraint terms. Now also for inhibitory weights (cf. Eq. 16):

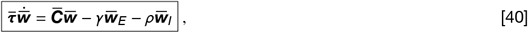

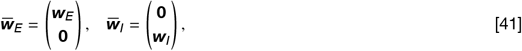

where **0** indicates vectors of zeros of appropriate dimension (*N*_*I*_ and *N*_*E*_) that we do not specify for better readability.

The constraint factors *γ* and *ρ* follow from the requirement that the weight vector does not grow along the direction of the constraint vectors 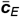 and 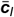. Here we choose them such that the sums over the excitatory and inhibitory weights remain constant, i.e., the L1-norm of the excitatory and inhibitory part of the weight vector is maintained^2^.

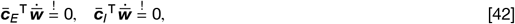

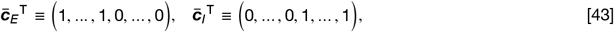

where the number of non-zero entries in 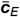 and 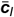 is equal to the number of excitatory *N*_*E*_ and inhibitory neurons *N*_*I*_, respectively. Based on these requirements we derive expressions for the scalar constraint factors *γ* and *ρ*:

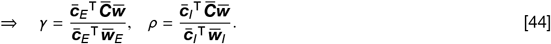

Finally, we can write the weight dynamics as

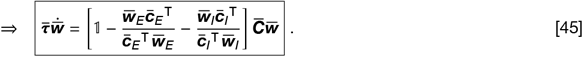

#### 2.1 Fixed points

For the fixed points we have to find weight vectors 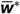 for which the time derivative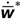 is equal to zero:

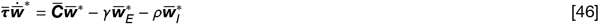

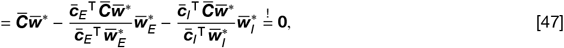

which is equivalent to

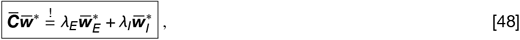

for *λ*_*E*_ and *λ*_*I*_ being arbitrary scalar.

##### 2.1.1 Eigenvectors of the modified covariance matrix are fixed points

It is straightforward to check that multiples of eigenvectors 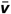 of the modified covariance matrix 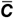 with eigenvalue 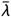 are fixed points:

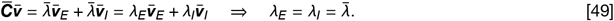

In the following, we will refer to eigenvectors of the modified covariance matrix as fixed point eigenvectors, and to eigenvectors of the feedforward excitatory covariance matrix ***C*** as feedforward eigenvectors. This leaves us with the task of finding the eigenvectors of 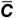. In general, eigenvectors depend non-trivially on the tuning of the laterally projecting population (cf. Sec. 3, Eq. 121). However, the problem simplifies when the laterally projecting inhibitory neurons are tuned to multiples of eigenvectors of the excitatory population’s covariance matrix. This is what one would expect when the post-synaptic excitatory neuron *r* and the inhibitory population ***y***_*E*_ both receive excitatory input from the same external brain region ***y***_*E*_ and synapses from the external population onto inhibitory neurons are plastic according to a Hebbian rule with multiplicative normalization (cf. Fig. S1*C*). Although we showed in Section 1.2.2 that without recurrent interactions only the principal eigenvector is a stable fixed point, we will find that with suitable recurrent interactions any feedforward eigenvector can be stable (cf. Sec. 3 & 5.2.3). Formally we set

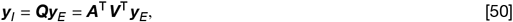

where each row of ***Q*** = ***A***^⊤^***V***^⊤^ is the feedforward weight vector of an inhibitory neuron which is equal to a positive multiple, *a*, of an eigenvector ***v*** of the excitatory covariance matrix ***C*** = ⟨***y***_***E***_ ***y***_***E***_ ^⊤^ ⟩. Then ***V*** holds all eigenvectors as columns, and ***A*** is a matrix where each multiple is the only non-zero element per column, such that ***AA***^⊤^ is a diagonal matrix. We will now show that in this scenario multiples of the excitatory and inhibitory part of the eigenvectors of the modified covariance matrix 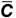 are fixed points. As a first step, we explicitly calculate the eigenvectors.

##### 2.1.2 Eigenvectors and eigenvalues of the modified covariance matrix

In the previous section, we have seen that eigenvectors 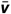 of the modified covariance matrix 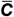 are fixed points. In this section, we will find an explicit expression for these eigenvectors when inhibitory neurons are tuned to feedforward eigenvectors, i.e., inhibitory neurons are tuned to eigenvectors ***v*** of the excitatory covariance matrix ***C***. Making use of Eq. 50 the modified covariance matrix becomes

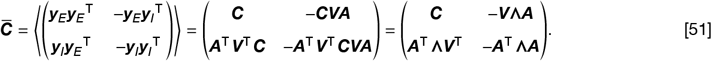

Then, a full set^1^ of linearly independent eigenvectors 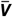 and their inverse 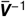 is given as^2^.

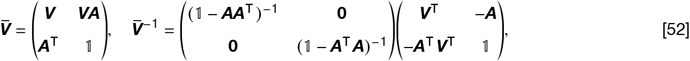

where each column of 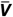 is an non-normalized eigenvector. The eigenvalue spectrum is

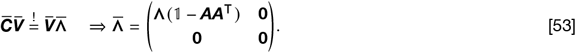

Similar to before, we call eigenvectors of the modified covariance matrix with positive eigenvalue *attractive*. Different from the case of only excitatory feedforward input, eigenvalues of the modified covariance matrix can also be negative. In this case, we call the corresponding eigenvector *repulsive* (cf. Sec. 1.1).

For eigenvectors in the right matrix column of 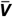 in Eq. 52, the excitatory and inhibitory components of the membrane potential exactly cancel, post-synaptic firing rates are zero, and no plasticity is induced: For multiple post-synaptic neurons with firing rates ***r***, where each neuron is tuned to one of these eigenvectors, one gets

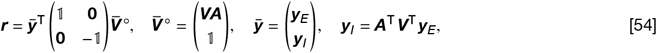

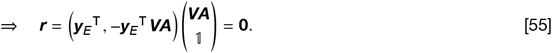

Since these eigenvectors result in post-synaptic firing rates of zero, and they define the null space of the 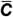 matrix (Eq. 53), we call them ‘null eigenvectors’ or ‘null fixed points’, and all eigenvectors that are not null eigenvectors ‘regular’ eigenvectors or fixed points. Note that for each additional inhibitory neuron that is tuned to a feedforward eigenvector, there is an additional null eigenvector, since inhibitory synaptic weights can now shift between the original, and the additional inhibitory neuron to cancel post-synaptic firing. Overall, there are always *N*_*I*_ null eigenvectors and *N*_*E*_ regular eigenvectors^3^.

We have already shown in Section 2.1.1 that eigenvectors of 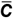 are fixed points. Each eigenvector specifies an exact ratio between the excitatory and inhibitory weight norm. Since our learning rule separately maintains the total excitatory and inhibitory synaptic weights, reaching any of these fixed points would require detailed fine-tuning at the point of initialization. In the next section, we show a more general set of fixed points that does not require any fine tuning of weight norms.

#### 2.1.3 Non-eigenvector fixed points

In this section, we show that there exist fixed points that are not eigenvectors of the modified covariance matrix. In particular, arbitrary multiples of the excitatory and inhibitory parts of regular eigenvectors, i.e., of eigenvectors that result in non-zero post-synaptic activity, are fixed points. We make the ansatz that the matrix 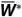 holds fixed points as columns and has the shape

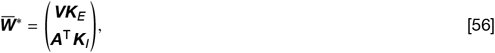

where ***K***_*E*_ and ***K***_*I*_ and are diagonal scaling matrices of arbitrary constants. The fixed point condition that follows from Eq. 48 is

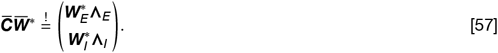

We now show that for any ***K***_*E*_, ***K***_*I*_ we can find diagonal matrices **Λ**_*E*_, **Λ**_*I*_ that fulfil this condition1. We write explicitly

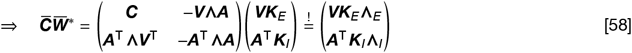

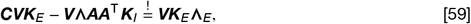

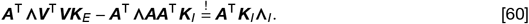

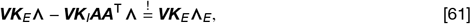

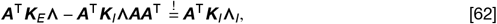

where we made use of the fact that independent of their subscript, the ***K*, Λ**, and ***AA***T commute. By comparing the left and right sides of the equations, we find

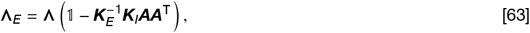

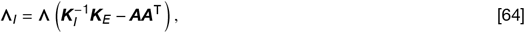

which are diagonal matrices, as required^2^. Before we consider the stability of these fixed points in Section 2.2, we first show that there is an additional set of fixed points.

##### 2.1.4 General fixed points

Having covered various special cases of fixed points for the dynamics, we now consider the general problem. Recall that fixed points are defined to satisfy Equation 48:

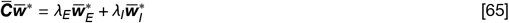

Expanding this using our expression for 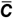 (Eq. 51), we can see that this is equivalent to:

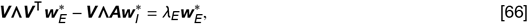

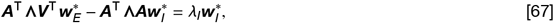

and equivalently

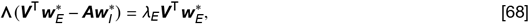

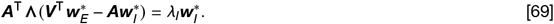

Inserting the first into the second expression, we can conclude that

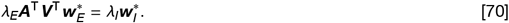

If *λ*_*E*_ = *λ*_*I*_ ≠ 0, then we know that 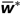 is an eigenvector of the modified covariance matrix, as discussed in Section 2.1.1. In the case that *λ*_*E*_ = *λ*_*I*_ = 0, we have the null eigenvectors discussed in Section 2.1.2. We therefore now address the case that *λ*_*E*_ ≠ *λ*_*I*_.

We begin with the case *λ*_*I*_ ≠ 0. Then we can insert Eq. 70 into Eq. 68 to arrive at

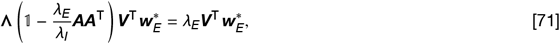

which, together with Eq. 70, gives necessary and s ufficient co)nditions for a fixed point. From Eq. 71, we conclude that 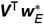 is an eigenvalue of the diagonal matrix 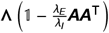 with eigenvalue *λ*_*E*_. When 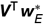 is one-hot, then the vector 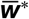consists of an arbitrary multiple of the excitatory and inhibitory parts of a regular eigenvector, as covered in Section 2.1.3.

We now turn our attention to the case where 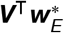 is not simply one-hot. We can now say that for each component *j* of 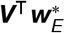 which is non-zero, the following equation must hold:

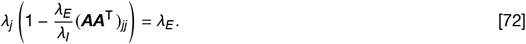

This is a linear system in the pair of variables *λ*_*E*_ and *λ*_*E*_ *λ*_*E*_. We work under the mild assumptions that the eigenvalues *λ*_*j*_, the diagonal elements (***AA***^⊤^) _*jj*_, and their product *λ*_*j*_ (***AA***^⊤^) _*jj*_ are distinct for each *j*. These conditions will in general hold in the absence of fine tuning. In this case, *λ*_*E*_ and *λ*_*I*_ provide two degrees of freedom and there will only be solutions when 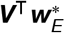 is (at most) two-hot, having non-zero components, *j* and *k*. Such solutions satisfy:

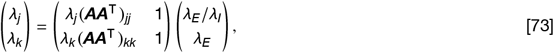

which we can solve to obtain the expressions:

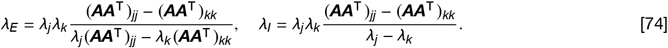

The components of the two-hot solution are determined by the initial values of ***c***_*I*_ ^⊤^***w***_*I*_ and ***c***_*E*_ ^⊤^***w***_*E*_, which are kept constant throughout training. Although two-hot fixed points do not require fine tuning of excitatory and inhibitory weight norms, we did not observe them in any of our numerical simulations and therefore assume they are unstable.

The final case to be considered is when *λ*_*I*_ = 0, *λ*_*E*_ ≠0. In this situation, Eq. 70 tells us that 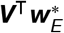 is in the kernel of ***A***^⊤^ and therefore in the kernel of the diagonal matrix ***AA***^⊤^1. By using Eq. 68, we can therefore conclude that

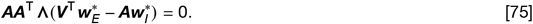

We work under the assumption that, in the absence of fine tuning, **Λ** has distinct non-zero eigenvalues. In this case, the first term in Equation 75 is zero, and 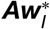 and therefore in the kernel of ***AA***^⊤^. So 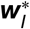 is in the kernel of ***A***^⊤^***A*** and therefore the kernel of ***A***. By Equation 68, this tells us that 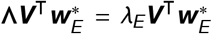 and therefore 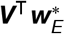 is an eigenvector of **Λ** with eigenvalue *λ*_*E*_. We therefore arrive at a fixed point for the system in which 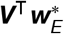 is one-hot with support on the kernel of ***AA***^⊤^, and 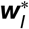is in the kernel of ***A***. This implies 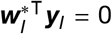 Eq. 50) which is biologically implausible since we constrain synaptic weights 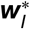 and firing rates ***y***_*I*_ to be positive.

Under mild assumptions regarding **Λ** and ***AA***^⊤^, we have thus exhaustively characterized the fixed points of the system.

### 2.2 Stability analysis

We first consider the stability of fixed points that are regular eigenvectors of the modified covariance matrix and discuss the case of non-eigenvector fixed points afterwards. With Eq. 45, for the Jacobian ***J*** it follows (cf. Eq. 29)

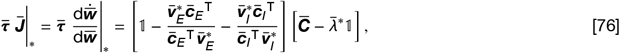

where 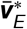 and 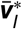 are the excitatory and the inhibitory part of the eigenvector fixed point 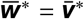 with eigenvalue 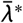, with an additional set of zeros to reach the correct dimensionality of the vector (cf. Eq. 41). To find the eigenvalues 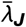of the Jacobian, we switch to the eigenbasis of the modified covariance matrix^2^:

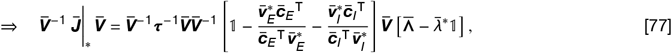

where we inserted 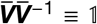. The result is a block triangular matrix where each block on the diagonal corresponds to one regular eigenvector and its potentially multiple null eigenvectors. To better see this, we consider the first and second part of Eq. 77 separately. We define ***e τ*** ^−1^, which remains a diagonal matrix with time constants for excitatory and inhibitory synapses on the diagonal, ***ϵ***_*E*_ = 𝟙, ϵ_*E*_ and ϵ_*I*_ = 𝟙, ϵ_*I*_. Inserting the definition of the eigenvectors matrix and its inverse (Eq. 52) we write

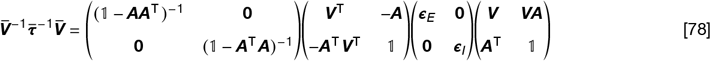

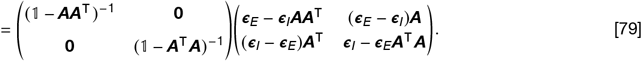

As one would expect, for ε_*E*_ = ε_*I*_, this is equal to a scalar times the identity matrix. When we switch columns and rows such that pairs of regular and corresponding null eigenvectors form blocks, this becomes a block diagonal matrix. Note that this does not change the determinant or the eigenvalues of the matrix as for each row switch, there is a corresponding column switch that maintains the characteristic polynomial. Alternatively, we can assume that the matrix of eigenvectors 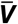 and its inverse 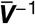 are already appropriately sorted. Without loss of generality, we assume that the first columns of 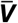 are the fixed point’s eigenvector 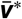 and its corresponding null eigenvectors, and write^1^

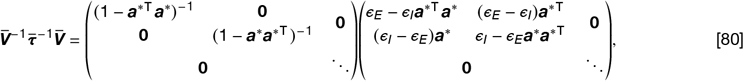

where ***a***^*^ is a column vector that holds the multiples of the inhibitory neurons that are tuned to the feedforward eigenvector ***v***^*^. As before, **0** are matrices of zeros and appropriate dimensionality, and ellipsis indicate continuing blocks on the diagonal with similar terms that belong to the non-fixed point eigenvectors and their null eigenvectors^2^.

Similarly, we can write the second part of Eq. 77 as a block triangular matrix. Before sorting, we write

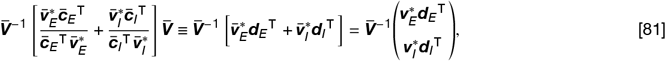

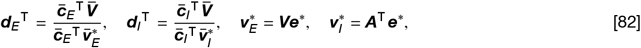

where ***d***_*E*_^⊤^ and *d*_*I*_^⊤^ are row vectors that hold the L1-norms of the eigenvectors’ excitatory and inhibitory parts as a fraction of the L1-norm of the fixed point eigenvector’s excitatory and inhibitory parts. The vector ***e***^*^ is zero except for one entry, equal to one, which corresponds to the fixed point feedforward eigenvector ***v***^*^. We continue by multiplying the inverse eigenvector matrix 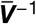 from the left:

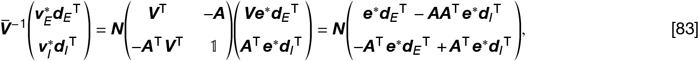

where we defined the normalization matrix ***N*** of the inverse eigenvector matrix 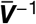 (cf. Eq. 52) to improve readability. It follows that the matrix above holds non-zero values in only a few rows, corresponding to the fixed point eigenvector (top block) and its null eigenvectors (bottom block). After rearranging, we get

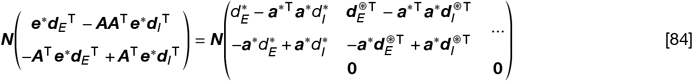

where 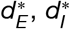 and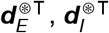 are the entries of *d*_*E*_^⊤^ *d*_*I*_^⊤^hat correspond to the fixed point eigenvector and its null eigenvectors, respectively. As before, ellipsis indicate additional non-zero entries. To find the respective entries of ***d***_*E*_, ***d***_*I*_ we use the definition of 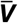 (Eq. 52) to write

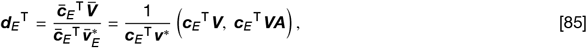

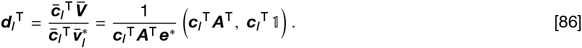

After rearranging the entries that correspond to the fixed point eigenvector and its null eigenvectors to the front we get

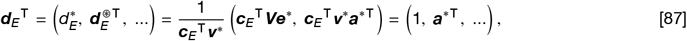

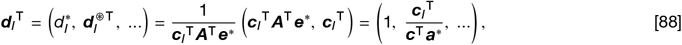

where ***e***^*^ selects the proper columns and ***c***^⊤^ is a row vector of ones of appropriate dimensionality. We insert Eq. 87 & 88 into Eq. 84 and find

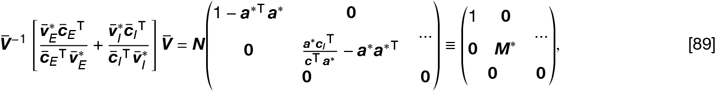

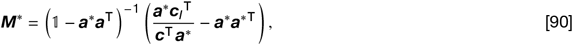

where we defined the matrix ***M***^*^.

In summary, we find that after rearrangement, Eq. 77 is a block triangular matrix.

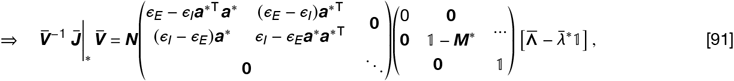

where we used Eq. 80 and Eq. 89. Therefore, to find the eigenvalues, we consider each diagonal block separately. We make the simplifying assumption that there is exactly one inhibitory neuron tuned to each feedforward eigen-vector. Then, ***a***^*^→ *a*^*^ becomes a scalar, ***N*** and ***A*** = ***A***^⊤^ become diagonal, and ***M***^*^ →1. The transformed Jacobian remains triangular and becomes

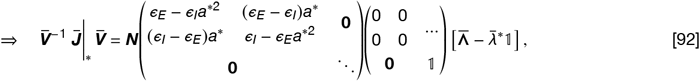

with 2× 2 blocks on the diagonal of which we only show the first, that corresponds to perturbations in the direction of the fixed point eigenvector or its null eigenvector^1^. From the matrix product above, we see that their corresponding eigenvalues must be zero since the first two columns of the second to last matrix are zero. For perturbations in the direction of a non-fixed point eigenvector 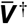 or its null eigenvector we have to consider the block matrix

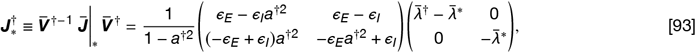

where 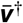 is a two-column matrix that holds 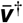 and its null eigenvector. The eigenvalues of this matrix are negative under two conditions. First, its determinant must be positive, and second, its trace must be negative. For trace and determinant, we find

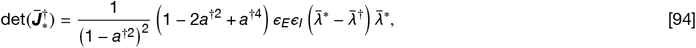

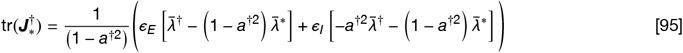

Finally, the two stability conditions read

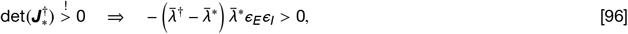

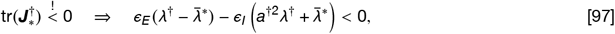

where, for the trace term, we made use of the equality 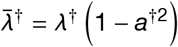 (cf. Eq. 53) to replace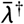.

#### 2.2.1 Principal component analysis in inhibition modified input space

The first stability condition above states that only the fixed point 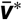 with the largest eigenvalue, 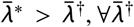, can be stable, and then only if it is not repulsive, i.e., provided that its corresponding eigenvalue is larger than zero. An eigenvector can become repulsive if inhibition is sufficiently strong, i.e., if 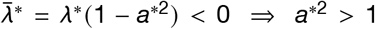. This implies that post-synaptic neurons tuned to repulsive eigenvectors receive more inhibition than excitation, which results in negative firing rates 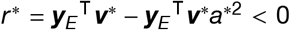 for 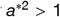 (cf. Eq. 55). However, in biology, neurons with larger inhibitory than excitatory input are hyperpolarized and remain silent, which is why we assume 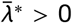. In the following, we call the combination of the excitatory feedforward attraction of an eigenvector *λ*^*^ (cf. Sec. 1.1) plus any contribution of laterally projecting neurons, in this case, minus the lateral inhibitory repulsion *a*^*2^*λ*^*^, the effective attraction, 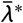, of a feedforward input mode.

For ε_*I*_ = ε_*E*_, the second condition reduces to^1^ 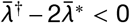, which holds if the first condition is met. Therefore, the post-synaptic neuron becomes tuned to the eigenvector of the modified covariance matrix with the largest eigen-value, i.e., it performs principal component analysis on a modified feedforward input space, where the attraction of feedforward eigenvectors is modified by laterally projecting inhibitory neurons (cf. Eq. 53). We will further discuss the notion of a modified input space in Section 3.

#### 2.2.3 Fast inhibition increases stability

In our networks, stationary states can still emerge when inhibitory plasticity is slower than excitatory plasticity. In the extreme case of static inhibition, ε_*I*_ = 0, the second stability condition is still satisfied if the fixed point attraction 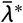 is larger than the feedforward attraction *λ*^†^ of any other eigenvector, 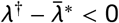. When inhibitory weights are static, they remain tuned to the fixed point and the repulsive component of competing eigenvectors *a*^†2^*λ*^†^ do not matter for stability. This explains why we have to consider only the attractive part *λ*^†^ of the effective attraction 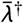 in the first term of the second stability condition. However, for growing *e*_*I*_ > 0, the influence of the inhibitory part of competing eigenvectors increases, corresponding to an increasingly negative second term in the second stability condition^2^. Then, for sufficiently fast inhibitory plasticity ε_*I*_ > ε_*E*_, the second condition always holds. Therefore, we consider slightly faster inhibitory than excitatory plasticity in our numerical simulations (cf. Table 1).

#### 2.2.3 Stability of non-eigenvector fixed points

Before, we considered the stability of fixed points 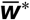 that are eigenvectors 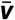 of the modified covariance matrix 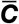. Weight vectors of that shape put a strong constraint on the choice of the weight norms, as the ratio between the excitatory and the inhibitory weight norms is given by the norms of the excitatory and the inhibitory parts of the eigenvector (cf. Eq. 52). The issue was solved in that we found that arbitrary combinations of multiples of the excitatory and inhibitory components of regular eigenvectors are also fixed points (cf. Sec. 2.1.3). We will now consider the stability of such non-eigenvector fixed points.

Let the shape of a fixed point 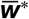 be (cf. Eq. 56)

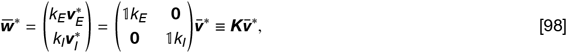

where *k*_*E*_ and *k*_*I*_ are scalar constants. We recapitulate the general weight dynamics as given in Eq. 45:

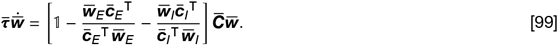

Instead of evaluating the eigenvalues of the Jacobian, we now switch to a new coordinate system in which the Jacobian will have a familiar shape. This is possible since fixed points and their stability do not depend on the choice of coordinates. We define:

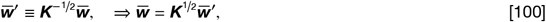

from which the weight dynamics can be written as

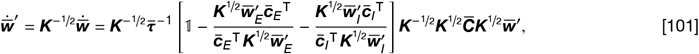

where we inserted ***K***^−1/2^***K***^1/2^ = 𝟙,. We now make use of the following identities:

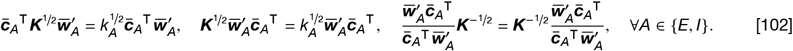

We find that the ***K***^1/2^ matrices inside the bracket cancel, and we can pull ***K***^−1/2^ from the right side to the left side of the bracket:

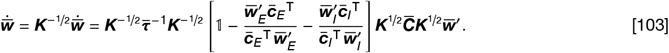

We introduce the following definitions

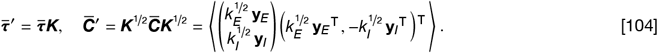

Note that 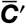 is *not* the modified covariance matrix expressed in the new coordinate system but a new modified covariance matrix that corresponds to an altered input space where excitatory and inhibitory input firing rates ***y***_*E*_, ***y***_*I*_ are scale by 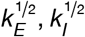, respectively. In summary, we can write the plasticity of the weight vector in the new coordinate system as^1^

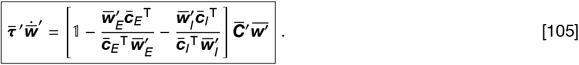

We are interested in the stability of the fixed points given in Eq. 98. In the new coordinate system, they become

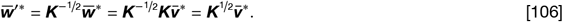

It is straightforward to proof that 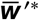 is an eigenvector of the new modified covariance matrix 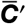 with eigenvalue 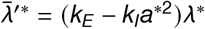: With 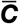 defined in Eq. 51 we get

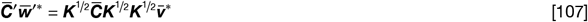

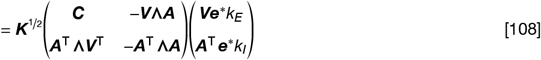

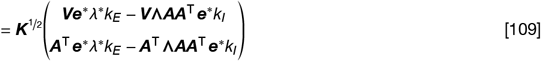

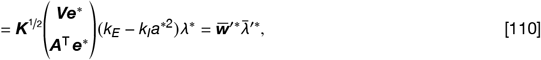

where we defined *a*^*2^ as the entry of the diagonal matrix ***AA***^⊤^ that corresponds to the eigenvector ***v***^*1^. are a regular eigenvectors of a new modified covariance matrix 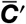. Note that this is independent of the change of variables, however, only in the new coordinate system one can identify the new modified covariance matrix with an actual input space^2^, where pre-synaptic firing rates are scaled by *k*_*E*_, *k*_*I*_ (Eq. 104). In theory, we can now proceed in finding the eigenvalues of the Jacobian^3^, as explained in Section 2.2. As before, one finds that stability is largely determined by the eigenvalues of the modified covariance matrix, which now are 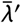.

Apart from providing a principled way to determine if a non-eigenvector fixed point is stable, our formulation provides additional insight: Let’s assume the total synaptic inhibitory weight of a neuron is very small, much smaller than any eigenvector of 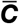 would suggest, i.e., *k*_*I*_ ≪ 1, while the excitatory weight norm is equal to one, which implies *k*_*E*_ = 1. As one would expect intuitively, the neuron does not exhibit much of the repulsion of the inhibitory neurons (cf. Eq. 104 for *k*_*I*_ ≪ 1), and its stability would be primarily determined by the excitatory attraction of the different eigenvector modes, i.e., 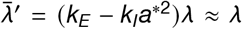. In the extreme case, when the inhibitory weight norm is zero, i.e., *k*_*I*_ = 0, only the activity of the excitatory population is relevant.

While the effective plasticity timescale 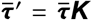 in Eq. 105 depends on the magnitude of the excitatory and the inhibitory part of the specific fixed point under consideration, this does *not* mean that the speed of synaptic plasticity is different from the original formulation in Eq. 45. For example, when we consider a fixed point with a decreased inhibitory weight norm *k*_*I*_ < 1, the effective inhibitory plasticity appears to increase, since 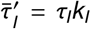. However, this effect is balanced by the decrease in pre-synaptic inhibitory firing rates, which decreases with decreasing *k*_*I*_^4^. Similarly, the coordinate system in which we describe the weight dynamics also does not affect the speed of plasticity^5^.

### 3 Lateral input stretches and compresses the feedforward input space

Before we consider how synapse-type-specific Hebbian plasticity affects learning in fully plastic recurrent networks, we first build additional intuition for how static lateral input affects the weight dynamics. From the previous section we know that in this case the eigenvalues of the modified covariance matrix are the key factors that determine fixed point stability, and from Sections 1.1 & 2.1.2 we know that these eigenvalues describe the Hebbian growth towards the corresponding eigenvector that can be attractive or repulsive, corresponding to a positive or negative eigenvalue. When a neuron receives only feedforward excitatory input (Fig. S2*A*), the weight dynamics is described by a true covariance matrix with eigenvalues equal to the variances along the principal components of the feedforward input space (cf. Sec. 1). Then the weight vector in the fixed point aligns with the direction of maximal variance in the input space (Fig.1*G*). In the following, we introduce a similar perspective and show that additional lateral input can be interpreted to stretch and compress the original feedforward input space, while the feedforward component of the weight vector performs PCA on this modified input space.

**Figure S2:**
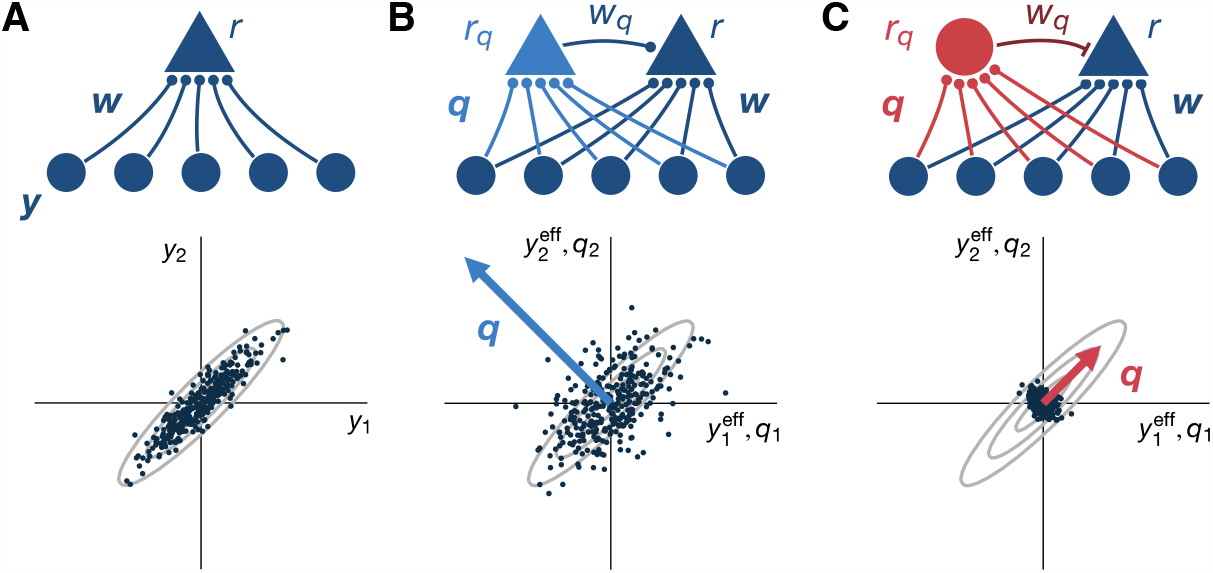
Input space modification due to lateral input. (*A*) Top: a single neuron with firing rate *r* receives synaptic inputs ***w*** from a population of excitatory neurons ***y***. Bottom: input distribution projected onto the first two input dimensions. Each dot represents the firing rates of the first two neurons during one input pattern. (Contour lines in light gray). Under a linear Hebbian learning rule, the neuron becomes selective for the direction of maximum variance, the first principal component (cf. Sec. 1). (*B*) Top: Same as in *A* for a neuron that receives additional input *w*_*q*_ from a laterally projecting excitatory neuron *r*_*q*_ which is tuned to an eigenvector ***q*** of the original input covariance matrix. Bottom: the effective input space ***y***^eff^ of the target neuron (dark blue triangle) is warped such that the variance along the eigenvector ***q*** (blue arrow) is stretched in proportion to the absolute value of the weight vector ***q***. The contour lines of the original input distribution from *A* are shown in light gray for reference. (*C*) Top: Same as *B* for a laterally projecting inhibitory neuron. Bottom: Now, the effective input space is compressed. See text for details.

We consider a circuit of two neurons that both receive feedforward input from a population of input neurons ***y*** (Fig. S2*B*, top). Let the first neuron have a fixed, non-plastic set of feedforward weights ***q*** and firing rate

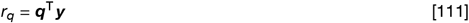

We let the first neuron project laterally onto the second neuron via a synaptic weight *w*_*q*_, without receiving any lateral input itself. Then the equilibrium firing rate of the second neuron is

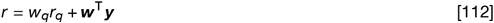

where we assume that both ***w*** and *w*_*q*_ are plastic according to a stabilized Hebbian rule.

From the perspective of the second neuron, the input space is increased by one dimension due to the additional lateral input, i.e., we can write Eq. 112 as

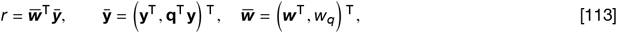

where, we defined the new input vector 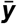 and the combined input weights 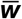. Effectively this is still a feedforward network without feedback, and the static covariance matrix 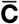 of the new inputs 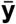 fully determines the average synaptic weight dynamics^1^:

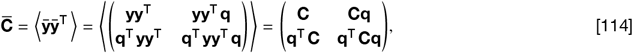

where ***C*** is the covariance matrix of the original input ***y***. We are interested in the eigenvectors 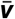 and eigenvalues 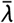 of this matrix for two reasons. First, because they describe the attraction towards different input modes due to the Hebbian term in our competitive plasticity rule. Second, because eigenvectors are fixed points with their stability mostly determined by the eigenvalues. Eigenvectors and eigenvalues must satisfy

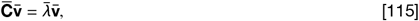

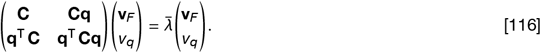

where *v*_*q*_ and ***v***_*F*_ are the lateral and the feedforward components of the eigenvector, respectively. In the following, we focus on the feedforward component for different ***q***. It follows

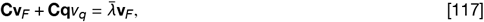

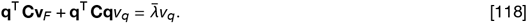

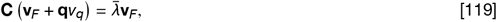

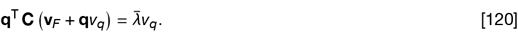

Inserting the first into the second expression gives *v*_*q*_ = **q**^⊤^**v**_*F*_ which, when inserted into the first expression, results in:

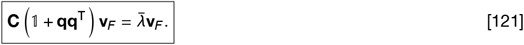

This is again an eigenvector equation, where the feedforward components of the original eigenvector ***v***_*F*_, are them-selves eigenvectors of the matrix **C(**𝟙,+ **qq**^⊤^ ). Note that for ***q*** = **0** we recover the case without lateral projections and feedforward components are multiples of eigenvectors of ***C*** with attractions 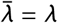. For general ***q***, the solution is not straightforward: We consider the equation in the input eigenspace, where Eq. 121 becomes

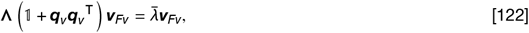

with **Λ** being the diagonal matrix of feedforward eigenvalues *λ* and the subscript (·) _*v*_ indicates a vector in the eigen-basis of ***C***. In this basis, eigenvectors of ***C*** are unit vectors, i.e., ***v***_*v*_ = ***e***, where ***e*** is a vector of zeros with a single entry equal to one, corresponding to the respective eigenvector. When ***q*** contains components of more than one eigenvector, the matrix ***q***_***v***_ ***q***_***v***_ ^⊤^ is not diagonal and eigenvectors of ***C***, do not solve the equation. Here we consider a simplified case: When the first neuron had plastic feedforward input, we know from Section 1 that it would converge to a multiple of an eigenvector of the feedforward covariance matrix^1^, ***q***∝ ***v***^†^, with ***Cv***^†^ = *λ*^†^***v***^†^. Then, ***q***_*v*_***q***_*v*_^⊤^e^†e†⊤^ is diagonal with a single non-zero entry, and it is obvious that feedforward eigenvector components of ***C*** are eigen-vectors of the feedforward covariance matrix ***C*** that solve Eq. 121.

To find the eigenvalues 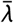, we first consider feedforward eigenvector components ***v***_*F*_ that are orthogonal to ***q***:

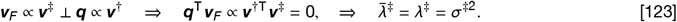

Therefore, input modes that are orthogonal to the tuning of the laterally projecting neuron maintain their attractions, equal to the respective eigenvalues of ***C*** that is equal to the variances *σ*^‡2^ of the input distribution along the respective orthogonal eigenvectors. Therefore, the laterally projecting neuron does not affect the Hebbian growth dynamics in the input subspace orthogonal to ***q***^2^. The remaining feedforward eigenvector component is proportional to ***q***:

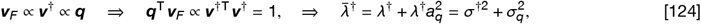

where *a*_*q*_ is equal to ∥***q***∥, the L2-norm of ***q***, and 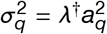 is the firing rate variance of the laterally projecting neuron. Therefore, the second neuron adjusts its feedforward weights ***w*** as if the variance along the eigenvector ***q*** was increased by 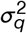 (Fig. S2*B*, bottom). In that sense, the second neuron ‘perceives’ its feedforward input space as stretched and we speak of a modified input space (cf. Sec. 2.2.1) that is described by a modified covariance matrix 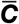. We note that it is possible to choose ***q*** such that an arbitrary direction of the input space becomes stable. For ***q*** = ***C***^−1^***h*** Eq. 121 is^3^

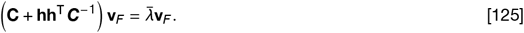

For increasing ∥***h***∥, the principal eigenvector transitions from 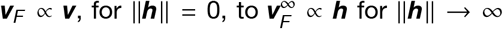. In the following, we only consider the case when ***q*** is parallel to one of the eigenvectors of ***C***. Then, for sufficiently large *a*_*q*_ and ***q*** ∝ ***v***^†^, an arbitrary non-principle eigenvector ***v***^†^ with attraction 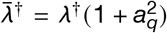 can become stable. In that case, the corresponding fixed point is of the following shape^1^:

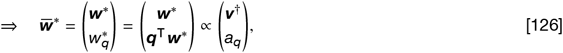

When the laterally projecting neuron is inhibitory (Fig. S2*C*, top), the modified covariance matrix becomes (cf. Eq. 51)

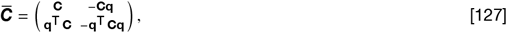

and it follows that the input space is compressed along ***q*** ∝ ***v***^†^ (Fig. S2*C*, bottom):

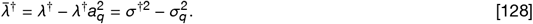

In the case of lateral inhibition and sufficiently large vector norms *a*_*q*_, an eigenvector can become repulsive, i.e., its eigenvalue becomes negative. Geometrically, this corresponds to a reflection of the input space along ***q*** through the origin, which can no longer be visualized as intuitively as in Fig. S2.

We can generalize the overall approach to multiple excitatory and inhibitory neurons such that the effective attraction towards a feedforward eigenvector becomes

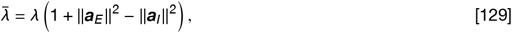

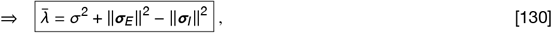

where *λ* = *σ*^2^, the vectors ***a***_*E*_, ***a***_*I*_ hold the feedforward vector norms of the laterally projecting neurons that are tuned to the respective feedforward eigenvector, and ***σ***_*A*_ = *λ****a***_*A*_, *A* ∈ { *E, I*}, hold the standard deviations of their firing rates. This allows writing the regular fixed points as^2^

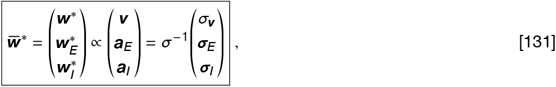

This implies that for regular fixed points, the total synaptic weight is distributed among synapses in proportion to the standard deviation of their pre-synaptic activities. Note that the different weight norms of the excitatory and inhibitory part of non-eigenvector fixed points can distort this relation (cf. Sec. 2.1.3).

In summary, we demonstrated how static lateral input can be interpreted to reshape the feedforward attraction landscape of afferent neurons. Note that these results are independent of what causes the laterally projecting neurons’ tuning. The second, afferent neuron does not ‘see’ what inputs to the laterally projecting neurons cause their tuning For example, in addition to feedforward input, laterally projecting neurons might be integrated into a recurrent circuit of neurons that are all tuned to the same eigenvector^3^. Then 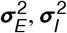 result from recurrent interaction in addition to the norm of the feedforward weight vectors. However, for the dynamics of the second neuron, it would not make any difference as long as the firing rate statistics of its pre-synaptic inputs were the same. In the following sections, we will consider circuits where the firing rate statistics emerge from recurrent interactions.

### 4 Eigencircuits

In the previous section we considered neurons that receive feedforward input from an excitatory population and lateral input from neurons with fixed feedforward tuning (Fig. S2). We found that the attraction of different feedforward input modes is determined by the eigenvalues of a modified covariance matrix, composed of a feedforward contribution and a contribution due to the laterally projecting neurons that is proportional to the variance of their firing rates (Eq. 130). In this section, we consider networks of recurrently connected, laterally projecting neurons and explore the variances of their firing rates.

First, we consider a network of excitatory and inhibitory neurons ***y***_*E*_, ***y***_*I*_ that are laterally connected to themselves and each other and receive feedforward input from the same excitatory population ***y***. We assume that the activity in the network is dominated by feedforward input such that neurons become selective for different eigenvectors of the feedforward covariance matrix ***C*** = ⟨***yy***^⊤^ ⟩, e.g., the steady state firing rate *y*_*a*_ of a neuron that is tuned to an eigenvector ***v***_*a*_ is proportional to ***v***_*a*_ ^⊤^***y*** (Fig. 4*A*), where the proportionality factor depends on the number and firing rates of other neurons that are tuned to the same eigenvector (see Sec. 4.1). Then the average Hebbian growth of a synapse that connects two neurons that are tuned to different eigenvectors is^1^:

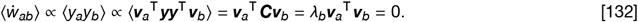

Due to the competition for synaptic resources, the synapse loses out to the non-zero growth of synapses that connect neurons that are tuned to the same eigenvector, and decays to zero over time (Fig. 4*B*). Eventually, the circuit is separated into sub-circuits that are tuned to different eigenvectors with recurrent connections within, but not between sub-circuits. Since there is one sub-circuit per eigenvector of the feedforward covariance matrix, we call these decoupled sub-circuits ‘eigencircuits’ (cf. Fig. 4).

#### 4.1 Variance propagation

In Section 3 we have seen that the attraction and the stability of a feedforward eigenvector is closely related to the firing rate variances of laterally projecting neurons, independent from how these variances arise. In the effective feedforward circuits that we considered, it was straightforward to compute variances based on feedforward weight norms (Eq.130 f.). We now show how variances can be determined in recurrent eigencircuits, which allows to quantify the effective attraction of an input mode.

We consider a generic eigencircuit and investigate how variances propagate through the network, i.e., our goal is to express the standard deviation of a neuron’s firing rate as a function of the standard deviations of its pre-synaptic input firing rates. For a neuron in an eigencircuit, all pre-synaptic inputs with non-zero synaptic weight are tuned to the same feedforward eigenvector ***v***. We only consider these non-zero entries and assume that the steady state firing rate of an arbitrary neuron can be written as (Fig. S3*A*)

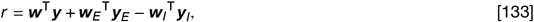

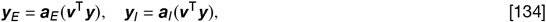

Note that before, ***a***_*E*_ and ***a***_*I*_ referred to feedforward weight norms (cf. Sec. 3). Now these vectors more generally express how firing rate variances relate to the input variance along the eigencircuit’s feedforward eigenvector, without making any assumptions about how this tuning arises. We will show in Section 5 that this assumption is correct and specify how the entries of ***a***_*E*_, ***a***_*I*_ relate to the recurrent excitatory and inhibitory weights (cf. Eqs. 160 & 161). For the weight vectors, we require that the excitatory and inhibitory parts are normalized to maintain the total amount of inhibitory and excitatory synaptic resources:

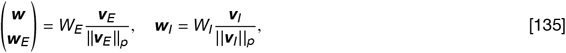

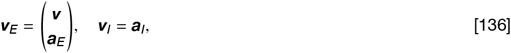

where *W*_*E*_, *W*_*I*_ are scalar weight norms, and ***v***_*E*_, ***v***_*I*_ are the excitatory and inhibitory parts of the fixed point eigenvector (cf. Sec. 5), with entries that are proportional to the pre-synaptic standard deviations (cf. Eq. 131). Then, the p-norm, ∥· ∥ _*p*_, is maintained due to competition for synaptic resources^2^. For the post-synaptic firing rate, it follows

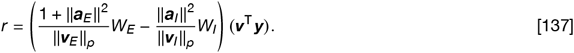

The first bracket is a scalar pre-factor which makes it straightforward to compute the standard deviation:

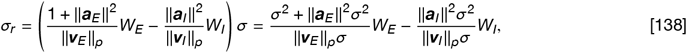

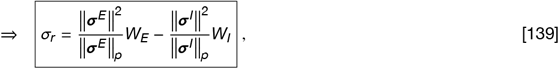

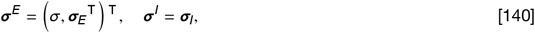

For a network in the steady state, i.e., when synaptic weights converged, this equation puts the standard deviation of neural firing rates in relation to each other, i.e., it provides the standard deviation of a post-synaptic neuron’s activity as a function of the standard deviations of its pre-synaptic input neurons’ activities^1^. It describes how standard deviations and variances ‘propagate’ through the network. In the next section, we will use this variance propagation equation (Eq. 139) to express the standard deviations in terms of only the weight norms and the feedforward standard deviation *σ*.

#### 4.2 Consistency conditions provide eigencircuit firing rate variances

We now consider a single eigencircuit where *n*_*E*_ excitatory and *n*_*I*_ inhibitory neurons are recurrently connected, and are tuned to the same feedforward eigenvector with standard deviation *σ* (Fig. S3*B*). In their steady state, all neurons have to fulfil the variance propagation equation (Eq. 139). In the fully connected eigencircuit, the firing rate variance of each neuron depends on the firing rate variances of all other neurons, and all neurons have the same set of non-zero pre-synaptic inputs. This provides *N* = *n*_*E*_ + *n*_*I*_ consistency conditions for the *N* unknown standard deviations. For example, the condition for a single excitatory neuron *i* reads

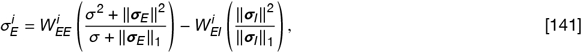

where we chose the L1-norm, *p* = 1, for normalization (but see Sec. 4.3), and *W*_*AB*_, *A, B E, I* are the total synaptic weight that a neuron of type *A* receives from neurons of type *B*. We make the simplifying assumption that all neurons have similar weight norms, i.e., 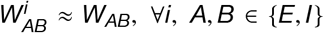. Then, also the standard deviations of their activities are similar, and we approximate 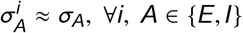

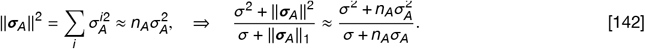

The standard deviations of excitatory and inhibitory neural firing rates become

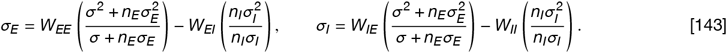

After some algebra, this yields the standard deviations as

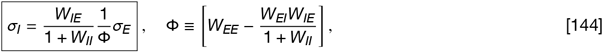

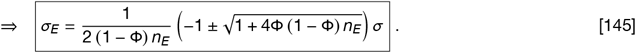

This provides standard deviations as a function of the number of neurons in the eigencircuit^2^, *n*_*E*_, *n*_*I*_, their weight norms, *W*_*AB*_, and the standard deviation of the feedforward input activity along the corresponding eigenvector, *σ*.Via Eq. 130, we can determine how the eigencircuit modifies the attraction of its feedforward eigenvector, i.e., the effective attraction 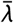 is

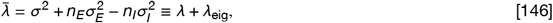

where we defined the attraction of the eigencircuit, *λ*_eig_, and *λ* is the attraction of the respective feedforward eigen-vector. In the following, we refer to 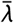 interchangeably as the effective attraction of the eigencircuit or the effective attraction of the feedforward input mode.

In summary, we assumed that neurons are tuned to feedforward eigenvectors (Eq.134) and showed how the network decomposes into recurrent eigencircuits. We demonstrated how variances propagate through such eigencircuits, and quantified how eigencircuits modify the attraction of their feedforward eigenvector (cf. Sec. 3) by laterally projecting onto other neurons (cf. Fig. 4*C*). In the following (Sec. 5), we will show that eigencircuits are indeed stable fixed points of fully plastic recurrent networks and investigate their stability.

#### 4.3 A note on the choice of weight norm

The choice of the weight norm that is maintained via multiplicative normalization is non-trivial. Biologically we motivated normalization by the competition for a limited amount of synaptic resources. We assumed the simplest case, where the L1-norm is maintained, and each resource unit translates to one unit of synaptic strength. An alternative choice would be to maintain the L2-norm. In the variance propagation equation (Eq. 139) this corresponds to *p* = 2 which becomes

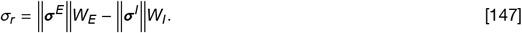

Following a similar logic as in Section 4.2, the eigencircuit consistency condition for a single inhibitory neuron becomes (cf. Eq. 141):

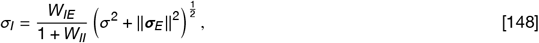

where we once more assumed that all neurons have similar weight norms, 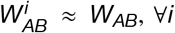. The variance of an excitatory neuron becomes

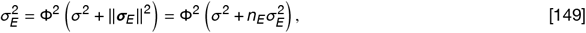

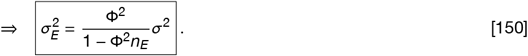

For an increasing number of excitatory neurons *n*_*E*_, the firing rate variance of a single excitatory neuron grows and diverges for Φ^2^*n*_*E*_ = 1. For even larger *n*_*E*_, variances would have to be negative to fulfil the consistency condition, which is not possible. It follows that for sufficiently large *n*_*E*_ there exist no fixed points. This is not unique to the L2-norm but holds for any *p* > 1. Such norms allow for a larger total synaptic weight (in terms of its L1-norm) when distributed across multiple synapses. Additional neurons provide additional recurrent synapses, which leads to the growth of the effective recurrent excitation until activities can no longer be stabilized by recurrent inhibition. For a suitable choice of the weight norms, Φ can, in principle, become small enough to balance the number of excitatory neurons in any eigencircuit to maintain positive variances. However, this requires additional fine-tuning and fails when *n*_*E*_ becomes unexpectedly large.

### 5 E-I networks with fully plastic recurrent connectivity

We now consider fully connected networks of excitatory and inhibitory neurons where all connections, feedforward and recurrent, are plastic according to the competitive Hebbian learning rule we introduced in Section 2. We will first show that eigencircuits are fixed points and then consider their stability with respect to a weight perturbation. Specifically, we would like to know when a neuron from one eigencircuit becomes attracted to a different feedforward eigenvector. We start with some simplifying assumptions.

**Figure S3:**
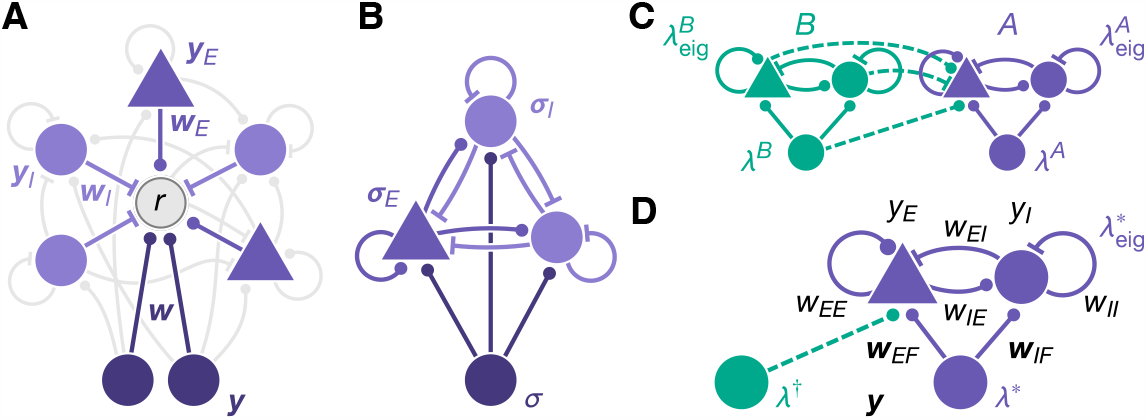
(*A*) A neuron with firing rate *r* (gray, center) receives synaptic inputs as part of a recurrent eigencircuit. The neuron receives excitatory synapses ***w*** from a population of input neurons ***y*** (dark purple, bottom). Excitatory (purple, triangles) and inhibitory neurons (light purple, circles) with firing rates, ***y***_***E***_, ***y***_*I*_, that are part of the same eigencircuit, project laterally onto neuron *r* via excitatory ***w***_*E*_ and inhibitory ***w***_*I*_ synapses. Recurrent synaptic connections that are not inputs of neuron *r* are shown in light gray – Not all synaptic connections are shown, for clarity. (*B*) Recurrently connected eigencircuit of *n*_*E*_ = 1 excitatory neuron (purple triangle) and *n*_*I*_ = 2 inhibitory neurons (light purple circles) that are tuned to the same feedforward eigenvector (dark purple circle, bottom). The standard deviation *σ* of input firing rates along the input eigenvector propagates through the network and results in firing rate standard deviations of ***σ***_*E*_ and ***σ***_*I*_ (cf. Eq. 144 f.). (*C*) Two excitatory neurons (triangles, top) and two inhibitory neurons (circles, top) in a recurrent circuit receive feedforward excitation from two input neurons (purple and green circles, bottom) that correspond to two different eigenvectors with eigenvalue *λ*^*A*^, *λ*^*B*^. Neurons are configured in a fixed point with two eigencircuits, *A* and *B*, with eigencircuit attractions 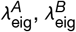 (cf. Eq. 146). Neurons that are part of the same eigencircuit are recurrently connected to each other. Synaptic weights between neurons that are tuned to different eigencircuits are zero. The excitatory neuron in eigencircuit *A* is perturbed in the direction of eigencircuit *B* (dashed lines). (*D*) Equivalent circuit with one excitatory and one inhibitory neuron. We consider a fixed point, where both neurons are tuned to the same feedforward eigenvector with eigenvalue *λ*^*^. The neurons form an eigencircuit with attraction 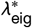. The excitatory neuron is perturbed in the direction of another feedforward eigenvector with attraction *λ*^†^ (dashed line). Firing rates, *y*_*E*_, *y*_*I*_, ***y***, and recurrent and feedforward synaptic weights, *w*_*EE*_, *w*_*EI*_, *w*_*IE*_, *w*_*II*_, ***w***_*EF*_, ***w***_*IF*_, are shown in black (cf. Eqs.152 & 152). See text for details.

Since each neuron can be bidirectionally connected to all other neurons, the dimensionality of the weight dynamics grows quadratically with the number of neurons. We are only interested in the general principles and consider a simplified circuit of two excitatory and two inhibitory neurons. One possible fixed point configuration is shown in Figure S3*C* (without dashed lines), where neurons are configured in two eigencircuits, *A,B*, with one excitatory and one inhibitory neuron per eigencircuit^1^. In this fixed point, all neurons receive feedforward input from a population of input neurons but synapses that connect neurons of different eigencircuits are zero (cf. Sec. 4). When a neuron in eigencircuit *A* is perturbed towards the other eigencircuit *B* (Fig. S3*C*, dashed lines), the tuning and the firing rates of all neurons in eigencircuit *A* change. However, neurons in eigencircuit *B* are unaffected because there are no connections projecting from eigencircuit *A* to eigencircuit *B*. Therefore, we only consider the recurrence within eigencircuit *A*, and think of input from other eigencircuits as effectively feedforward and static: That is, we construct an equivalent circuit where we perturb an excitatory neuron that is part of an eigencircuit, ‘^*^’, in the direction of another eigencircuit, ‘^†^’, that does not contain any neurons and has feedforward attraction equal to the effective attraction of eigencircuit *B*, that is^2^ (Fig. S3*D*)

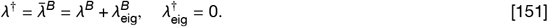

The configuration and attraction of the perturbed eigencircuit ‘^*^’ is equal to eigencircuit *A*, i.e., 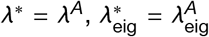. In Section 5.2.3 we will explain in more detail why these two circuits (Fig. S3*C & D*) are highly similar with regards to their stability.

In the equivalent circuit (Fig. S3*D*), we now consider the generic equilibrium firing rates of the *n*_*E*_ = 1 excitatory and *n*_*I*_ = 1 inhibitory neuron without taking any tuning into account (Fig. S3*D*)

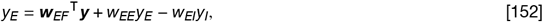

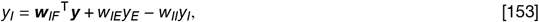

where ***y*** holds the firing rates of a population of *N*_*F*_ input neurons and we did not assume any specific tuning of the feedforward weights ***w***_*EF*_, ***w***_*IF*_. Since the network is linear, we can solve for the firing rates:

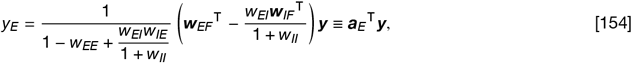

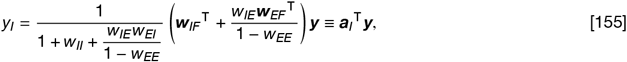

where we defined the effective feedforward vectors ***a***_*E*_, ***a***_*I*_. The weight dynamics is

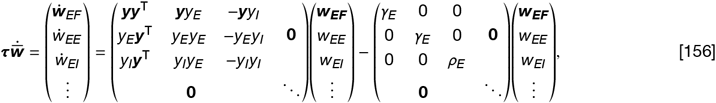

where ellipsis indicate similar terms for afferent weights of the inhibitory neuron. We define the modified covariance matrix

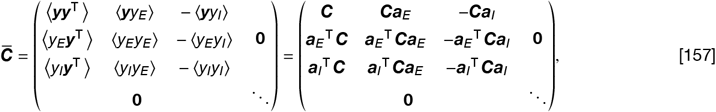

and write the average synaptic change as^1^ (cf. Eq. 40)

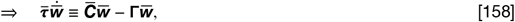

where **Γ** is a diagonal matrix that holds the scalar constraint factors, and 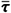 holds the timescales for excitatory synapses, ***τ***_*E*_ = 𝟙, *τ*_*E*_, and inhibitory synapses, ***τ***_*I*_ = 𝟙*τ*_*I*_, on the diagonal. We make the simplifying assumption that the plasticity of excitatory and inhibitory synapses is equally fast, *τ*_*E*_ = *τ*_*I*_ = *τ*. Then 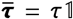, which does not affect the fixed points or the stability of the system^2^, and we set 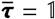.

Note that this is a highly non-linear dynamical system since the modified covariance matrix not only depends on the feedforward inputs ***y*** but also on the plastic synaptic weights 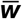, in addition to the weight dependence of the normalization factors **Γ**. Next, we show that the eigencircuit configuration we discussed in the introduction to this section is in fact a fixed point of the weight dynamics.

#### 5.1 Fixed points

In general, fixed points 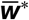 must fulfil the following condition

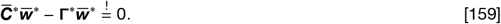

where 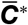 is the modified covariance matrix evaluated in the fixed point. We consider the special case when the two neurons form a single eigencircuit, tuned to the feedforward eigenvector ***v***^*^. Then we can write the excitatory and inhibitory firing rates as^3^

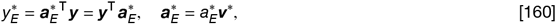

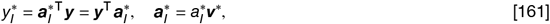

where *a*_*E*_ and *a*_*I*_ depend on the excitatory and inhibitory weights and can be determined via Eq. 154 & 155. This demonstrates that when neurons are tuned to the same feedforward eigenvector ***v***^*^, their firing rate is proportional to the projection of the activity vector ***y*** onto the eigenvector ***v***^*^, and justifies our assumption in Eq. 134. The modified covariance matrix in the fixed point becomes

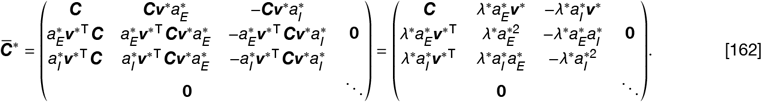

which can be diagonalized by the eigenvector matrix 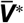 and its inverse:

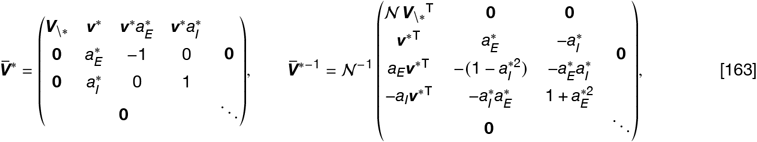

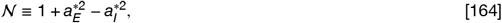

where the subscript (·)_\*_ indicates that a matrix does not contain an entry that corresponds to the input mode ***v***^*^.

In general, 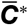 is of dimension 2*D*×2*D* where *D* = *N*_*F*_ +*N*_*E*_ +*N*_*I*_. Therefore, to diagonalize 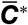, we require 2*D* eigenvectors. In this specific circuit, we have *N*_*E*_ = *N*_*I*_ = 1. Because 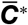 has a block diagonal structure (Eq. 162), with the first *D* ×*D* block driving development of weights onto the excitatory neuron and the second *D*×*D* block driving development of weights onto the inhibitory neuron, the eigenvector matrix 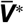 and its inverse have the same block diagonal structure. Since each block has the same sub-structure, we only show the first block in Eq. 163. Then for each eigencircuit there are 1 + *n*_*E*_ + *n*_*I*_ eigenvectors of 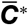, where *n*_*E*_ and *n*_*I*_ are the number of excitatory and inhibitory neurons in the respective eigencircuit: One regular eigenvector, and *n*_*E*_ +*n*_*I*_ null-eigenvectors (cf. Sec. 2.1.2). The first column of 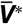 in Eq. 163 corresponds to the *N*_*F*_ − 1 eigencircuits with *n*_*E*_ = *n*_*I*_ = 0, i.e., with one regular eigenvector per feedforward eigenvector ***v*** ≠ ***v***^*^, but without null-eigenvectors. The corresponding eigenvalues are **Λ**_\*_, which are also eigenvalues of the feedforward covariance matrix ***C***. For the eigencircuit corresponding to ***v***^*^ there are 1 +*n*_*E*_+ *n*_*I*_ = 3 eigenvectors of 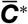. The first is a regular eigenvector and the last two are null eigenvectors, where the excitatory feedforward component is cancelled by either a negative lateral excitatory component^1^, or a positive lateral inhibitory component. The null eigenvectors have eigenvalues equal to zero, and the eigenvalue of the regular eigenvector is 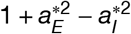.

Similar to the feedforward case, arbitrary multiples of the separately normalized parts of eigenvectors of 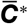 are fixed points. The only exception is the rightmost null eigenvector (cf. Sec. 2.1.3). There, the inhibitory and the excitatory weights are aligned such that the post-synaptic activity is zero, which does not allow for arbitrary scaling of the weight norms. Inserting these fixed points into Equations 154 & 155, provides conditions to determine 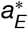 and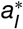

#### 5.2 Stability analysis

We are interested in the stability of the circuit described in the introduction of Section 5 and consider the stability of a regular eigenvector 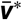

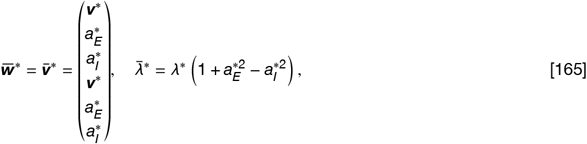

This means we do *not* consider arbitrary scalings of the excitatory and inhibitory parts of eigenvectors of 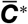, but assume that weight norms are fine tuned to match the norms of the excitatory and inhibitory parts of the regular eigenvector^1^ 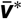.

When are such eigenvectors stable, and when are they attracted to a different input mode? To answer this question, we consider small fixed point perturbations 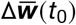, where the excitatory neuron shifts its tuning in the direction of a different feedforward input eigenvector ***v***^†^:

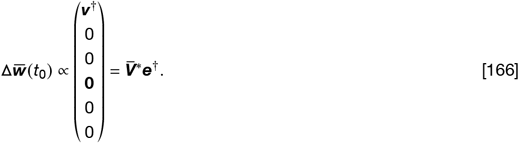

where ***e***^†^ is a vector of zeros with a single non-zero entry that corresponds to the feedforward eigenvector ***v***^†^ (cf. Eq. 163). The system is stable with respect to a perturbation if the perturbation decays to zero over time. To check this, we consider the following differential equation that holds for small perturbations (cf. Sec. 1.2.2)

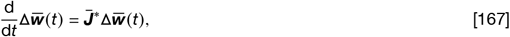

where 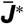 is the Jacobian evaluated in the fixed point. We will consider the dynamics in the non-orthogonal eigenbasis 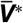 of the modified covariance matrix 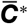 evaluated in the fixed point 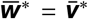. Note that 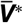 is not time-dependent, because it is evaluated in the fixed point. In this static basis, we can express perturbations as

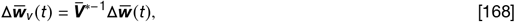

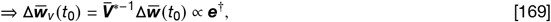

where the subscript (·)_*v*_ indicates a vector or matrix expressed in this basis. The perturbation dynamics becomes

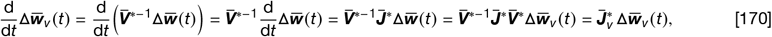

where we defined the transformed Jacobian, 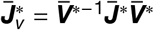 Without loss of generality, we assume that eigenvectors in 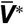 are sorted such that the first entry of ***e***^†^ is non-zero, i.e., the first column of 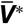 is proportional to the initial perturbation 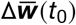 (cf. Eq. 166). Next, we will derive the transformed Jacobian.

##### 5.2.1 The transformed Jacobian

First, we consider the regular Jacobian 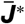. We rewrite the dynamics in Eq. 158 as^2^

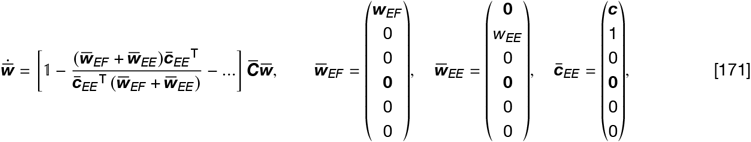

where the second term in the bracket corresponds to the normalization of all excitatory synapses onto the excitatory neuron, additional normalization terms are indicated by ellipsis^3^ (cf. Eq. 45), and ***c*** is a vector of ones. Then the Jacobian has the following shape (cf. Eq. 29)

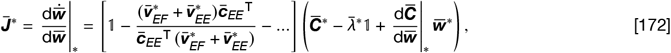

where 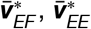 have the same shape as 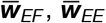 in Eq. 171 with entries corresponding to the respective entries of the regular eigenvector 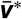 (cf. Eq. 165). Note that we accounted for the weight dependence of the modified covariance matrix 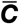 which results in the tensor 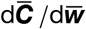. To find the transformed Jacobian 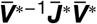, we consider the first bracket:

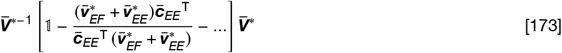

The first entry remains equal to the identity matrix, as the eigenvector matrix and its inverse cancel. We consider the columns 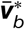 of 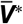 separately. Then, we can write

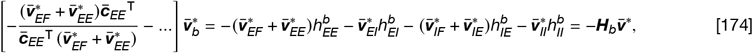

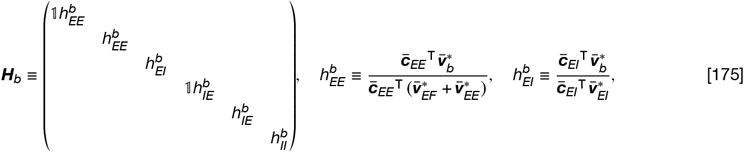

where ***H***_*b*_ is a diagonal matrix with entries corresponding to the respective normalization constraint, of which we give 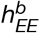 and 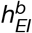 as examples. Then each column 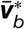 of 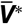 is transformed into a multiple of the separately normalized parts of the fixed point eigenvector 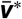 (Eq. 165). After transformation, the *b*th column becomes

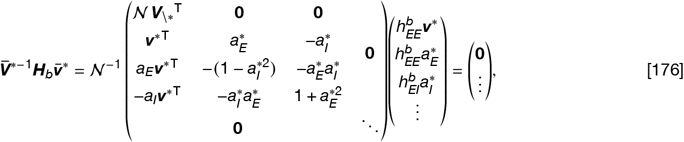

where, as before, ellipsis indicate potentially non-zero entries. Importantly, after the transformation, the first *N*_*F*_ − 1 entries are zero, independent of the column index, *b*, because ***v***^*^ is orthogonal to the columns of ***V***_\*_. Overall, we can write

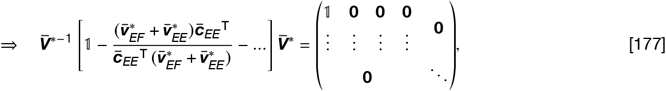

where the block structure arises from the block structure of 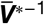(cf. Eq. 176).

After transformation, the second bracket of Eq. 172 becomes

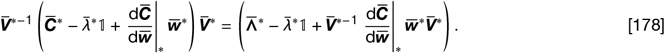

We next consider the first columns of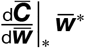, for which we compute the matrix 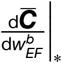, where 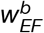 is the *b*th feed-forward weight onto the excitatory neuron.

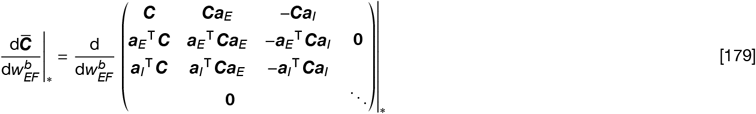

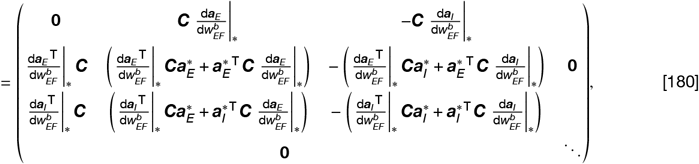

where we used the definition of 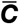 from Eq. 157. The vectors ***a***_*E*_ and ***a***_*I*_ are defined in Eq. 154 & 155. It follows:

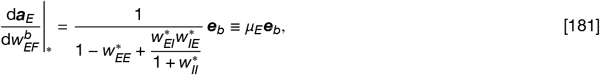

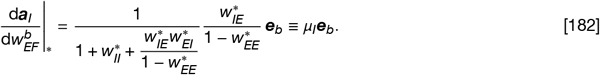

Additionally, we have (cf. Eqs.160 & 161) which results in

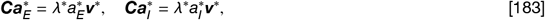

which results in

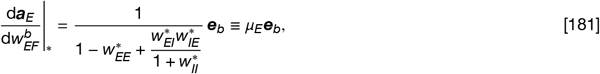

where ***e***_*b*_ is a vector of dimension *N*_*F*_ with entries equal to zero, except for the *b*th entry equal to one.

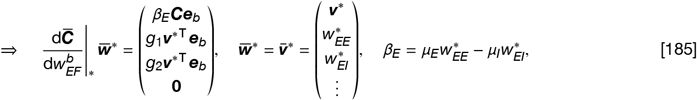

where *g*_(·)_ are scalars.

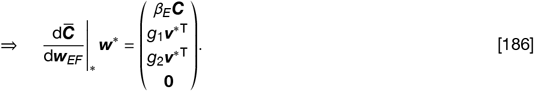

We find other columns in a similar fashion and write

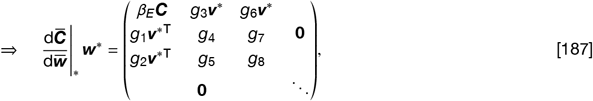

where, again, *g*_(·)_ are scalars. After applying the transformation, we get

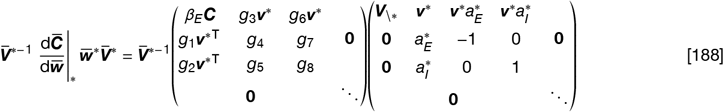

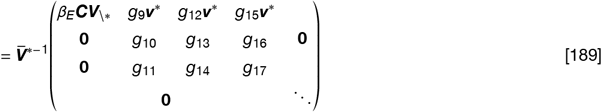

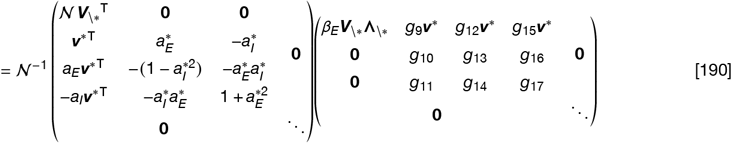

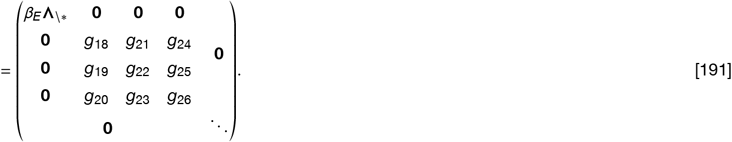

The fully transformed Jacobian is (cf. Eq. 178)

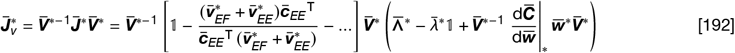

Finally, by inserting Eq. 177 & 191 we find

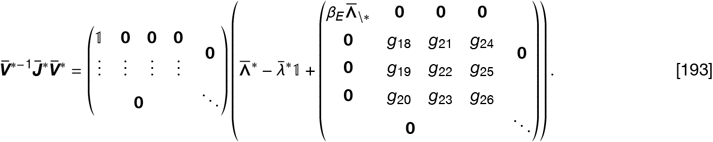

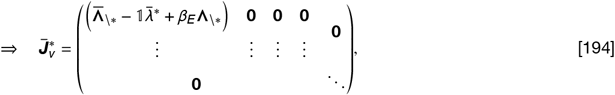

where 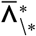 contains eigenvalues of 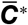 that correspond to regular, non-fixed point eigenvectors.^1^

##### 5.2.2 Stability conditions

The dynamics of a general fixed point perturbation Δ***w***_*v*_ in the eigenbasis of 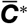 is (cf. Eq.170)

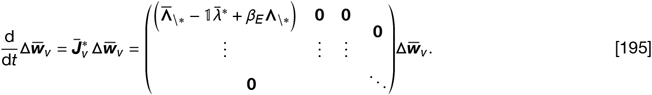

Note that the transformed Jacobian (Eq. 194) has a triangular block structure, and each row of 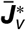 corresponds to the growth of a perturbation in the direction of a different eigenvector of 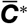. We are only interested in perturbations that grow in the direction of a non-fixed point feedforward eigenvector, ***V***\*. Therefore, we focus on the first rows of 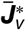, which correspond to growth in these directions. Except for the first diagonal block, these rows are zero. It follows that perturbations 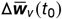 that do not already contain components in the direction of non-fixed point eigenvectors, also do not develop such components in their later dynamics. In contrast, perturbations in the direction of a non-fixed point feedforward input mode, e.g., 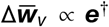, can induce perturbations within the original eigencircuit that corresponds to the feedforward eigenvector ***v***^*^ ^1^. For example, a decrease in feedforward and recurrent excitatory synaptic weights within the eigencircuit balances the increase of feedforward excitatory synaptic weights due to the perturbation towards a different eigencircuit, to maintain the weight norm. However, as explained above, these ‘second-order’ perturbations, without components in the direction of a non-fixed point feedforward eigenvectors, ***V***_\*_, are contained within the eigencircuit, i.e., they can not induce subsequent perturbations in the direction of non-fixed point feedforward input modes, ***V***_\*_. Therefore, to answer the question of when an eigencircuit is stable, we only consider the dynamics along the direction of the original perturbation (cf. Eq.166) by projecting the dynamics onto the perturbation vector at time zero, 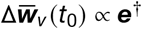 (cf. Eq.169):

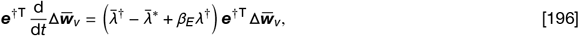

which provides the eigencircuit stability condition for the excitatory neuron

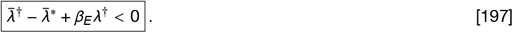

If Eq. 197 holds, perturbations in the direction of non-fixed point eigenvectors decay to zero, and the eigencircuit is stable. For *β*_*E*_ we have (cf. Eqs. 185 & 181 f.)

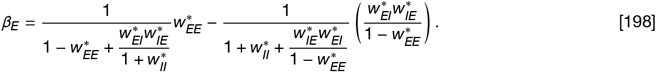

From Eq. 154 & 155 we find

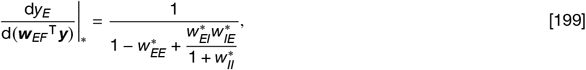

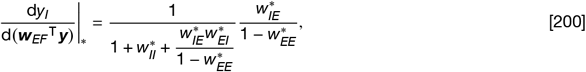

and we get

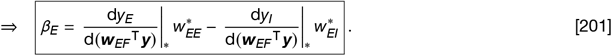

Following the same framework, we find the stability condition when perturbing the inhibitory neuron:

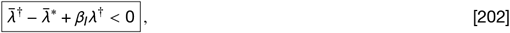

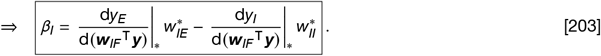

**Figure S4:**
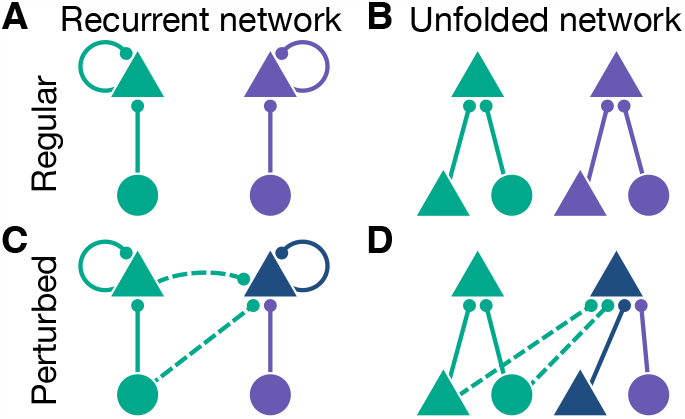
(*A*) Two excitatory neurons (triangles) are tuned to two different, but equally attractive input modes (circles, green and purple). (*B*) The same circuit as in *A*, unfolded to highlight pre-synaptic partners. Both input modes are balanced in their attraction. (*C*) Perturbing the purple excitatory neuron towards the green input mode (dashed lines) shifts its tuning (dark blue) such that it now responds to both the green and the purple input modes. (*D*) The unfolded circuit from *C*. Due to the perturbation, the green input mode is now more attractive, and the previously purple excitatory neuron shifts its tuning. See text for details.

We will now interpret these results.

##### 5.2.3 Eigencircuit stability depends on recurrent connectivity

We first consider the case when the effective attraction of all eigencircuits is the same, i.e., 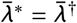 (cf. Eqs. 197 & 202). Then the stability is fully determined by *β*_*E*_, and *β*_*I*_. In feedforward circuits we have not found any *β* -terms, because in that case, the modified covariance matrix does not depend on any plastic synaptic weights (cf. Eq. 51 & Sec. 2.2). This is not the case in recurrent circuits where the perturbation induces a change in the tuning of laterally projecting neurons.

To build some intuition, we consider a simple example: Think of a recurrent network of two excitatory neurons with identical weight norms that receive feedforward input from the same population of excitatory neurons. In the fixed point, the neurons are tuned to two different feedforward input eigenvectors of equal attraction and are recurrently connected to themselves but not each other (Fig. S4*A*). Then the effective attraction of the two eigencircuits is the same. In general, neurons receive synaptic inputs, but have no information about the overall network structure, e.g., which synaptic inputs are feedforward or recurrent. Taking this perspective, we unfold the recurrent network and observe that the effective mode attraction is a combination of the feedforward input and the recurrent self-excitation (Fig. S4*B*). When we perturb one neuron towards the opposing input mode (Fig. S4*C*, dashed lines), the tuning of the perturbed neuron changes slightly in the direction of that mode (Fig. S4*C*, dark blue). From the perspective of the perturbed neuron, this tuning change leads to an attraction increase of the opposing eigencircuit, which is now more attractive than the original eigencircuit of the perturbed neuron (Fig. S4*D*), and the perturbation grows in the direction of the more attractive mode – the fixed point is unstable. Similarly, if the neurons were inhibitory instead, the perturbation would decrease the attraction towards the opposite input mode which would stabilize the network. In our mathematical analysis of the circuit shown in Figure S3*D*, the attraction increase or decrease due to the tuning change of recurrently projecting neurons is reflected in the *β* -terms in Equations 197 & 202, which emerge from the weight dependence of the modified covariance matrix 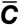 (cf. Eq. 172). For example, when perturbing the excitatory neuron, the increase in attraction from the perspective of the perturbed neuron is (cf. Eq. 197)

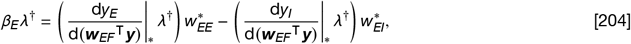

where the brackets reflect the tuning shifts of the excitatory and the inhibitory neuron^1^ in response to the perturbation of ***w***_*EF*_ in the direction of ***v***^†^, which is then weighted by the respective synaptic connection onto the excitatory neuron, 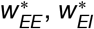,. When the inhibitory neuron is perturbed instead, the terms for *β*_*I*_ follow the same logic (cf. Eq. 203).

Without going through the lengthy mathematical derivation, we now give some intuition about *β* -terms of the network perturbation in Figure S3*C*. In the fixed point, the perturbed excitatory neuron receives recurrent input from all neurons in its eigencircuit, including itself. In the following, superscripts indicate the corresponding eigencircuit, *A* or *B*. Then, as for the equivalent circuit (cf. Fig. S3*D*), 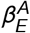 comprises two terms, one due to the tuning shift of 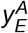, and a second due to the tuning shift of 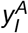. Assuming the same weight norms, this is exactly equal to the *β*_*E*_ for the equivalent circuit (Eq. 197). Different from 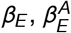is weighted with the effective attraction 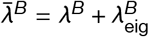, instead of only the feedforward attraction (cf. *λ*^†^ in Eq. 204), because the perturbation comprises not only the feedforward eigenvector component but the whole eigencircuit (cf. dashed lines in Figs. S3*C & D*). This is why, for the equivalent circuit, we chose the feedforward attraction 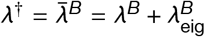 (Eq. 151). Then, the diagonal entries corresponding to the respective perturbations in the upper left blocks of the transformed Jacobians are the same^1^ (cf. Eq. 194), i.e.,

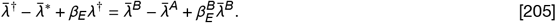

We find that perturbations in both circuits initially follow the same dynamics, while the later dynamics diverges: At time *t*_0_, there are no lateral projections from eigencircuit *B* towards eigencircuit *A* (cf. Fig. S3*C*), since in the fixed point there are no recurrent connections between eigencircuits (cf. Sec. 4), and the perturbation at time *t*_0_ only introduces connections from eigencircuit *B* onto eigencircuit *A*. However, as we just discussed, the perturbation introduces a tuning shift in neurons of eigencircuit *A* in the direction of eigencircuit *B*. This shift leads to non-zero correlations between neurons of both eigencircuits, and synaptic weights from eigencircuit *A* onto eigencircuit *B* start to grow. These growing synapses shift the attraction of neurons in eigencircuit *B* and thus impact the dynamics of perturbation components in the direction of eigencircuit *B*. Therefore, the transformed Jacobian of the original circuit (Fig. S3*C*) has a more complex structure than the Jacobian for the equivalent circuit^2^. However, since we consider an initial perturbation that is aligned with a regular eigenvector, i.e., 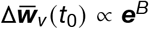 is one-hot (cf. Eq. 169), the top left diagonal block of the Jacobian still determines the initial dynamics^3^.

In summary, recurrent synapses can stabilize or destabilize a circuit with respect to small perturbations away from a fixed point. These stabilizing and destabilizing effects are described by *β* -terms that depend on the specific weight configuration in the fixed point (cf. Eq. 198), which again depends on the weight norms that constrain the total synaptic weights. In the following, we consider the case when synaptic weights are tuned such that *β* -terms are small. In the equivalent circuit (Fig. S3*D*) this is the case when the influence of the tuning shifts of the excitatory and the inhibitory neuron balance each other^4^ (cf. the first and second terms in Eqs. 201 & 203).

##### 5.2.4 Decorrelation condition

We now consider how neurons self-organize to represent all parts of their input space instead of clustering all their tuning curves around a dominant input mode. We consider the fixed point stability of different eigencircuit configurations. In particular, we consider the case when recurrent connectivity motifs do not influence the stability of an eigencircuit, i.e., *β*_*E*_ and *β*_*I*_ are approximately zero. This can be achieved by a suitable choice of weight norms^5^. Then, all eigencircuits are marginally stable when they are equally attractive, i.e., (cf. Eqs. 197 & 202 for *β*_*E*/*I*_ = 0)

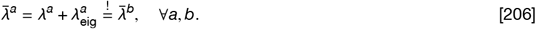

For homogeneous input spaces, where the feedforward attraction of all input modes is the same, i.e., *λ*^*a*^ = *λ*^*b*^ = *λ*, ⩝*a, b*, the only alternative stable configuration is when all neurons are tuned to the same feedforward input mode and form a single eigencircuit. Such a configuration does not reflect the tunings of biological neural populations, where all parts of the input space are represented. To prevent such a global clustering of neural tunings, we require that the corresponding eigencircuit is unstable. When all *β* -terms are approximately zero, this is the case when the effective attraction of the only occupied eigencircuit, 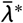, is smaller than the attraction of one of the *N*_*F*_ −1 unoccupied^6^ input modes, 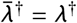 (cf. Eq. 197 & 202):

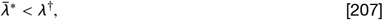

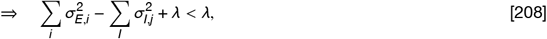

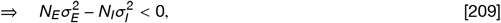

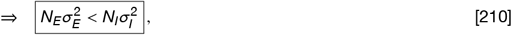

where 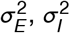 are the average variance, and *N*_*E*_, *N*_*I*_ the total number of inhibitory and excitatory neurons. When this condition is satisfied, the only stable solution is when the effective attraction of all eigencircuits is identical. The simplest configuration where this is the case is when each eigencircuit contains the same number of excitatory and inhibitory neurons.

### 6 Movie captions

**Movie M1: Decorrelation of feedforward tuning curves of excitatory neurons in plastic recurrent networks**. Development of feedforward tuning curves of *N*_*E*_ = 10 excitatory neurons (cf. Figs. 3*A & B*). Synaptic weights were initialized randomly. Different color shades indicate weights of different post-synaptic neurons.

**Movie M2: Decorrelation of feedforward tuning curves of inhibitory neurons in plastic recurrent networks**. Development of feedforward tuning curves of *N*_*I*_ = 10 inhibitory neurons (cf. Figs. 3*A & B*). Synaptic weights were initialized randomly. Different color shades indicate weights of different post-synaptic neurons.

## Footnotes

1 We indicate an equality or condition that we want to be fulfilled with an exclamation point over the equal sign.

1 This can be seen by formulating the system in the eigenbasis of ***J***^*^. Then, the matrix exponential becomes: ***V***^−1^ exp (***J***^*^) ***V*** = exp **Λ**_***J***_, where ***V*** holds eigenvectors and **Λ**_***J***_ is a diagonal matrix that holds the eigenvalues of ***J***^*^.

2 In general, the real part of the eigenvalues of the Jacobian have to be negative for a fixed point to be stable. However, since *C* is a covariance matrix, it is positive definite with positive, real eigenvalues. We will see (Eq. 32) that from this it follows that the eigenvalues of the Jacobian are also real.

1 To make sense of the vector notation, it helps to first consider the *b*’th column of 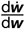 which is equal to 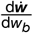, where *w*_*b*_ is the *b*’th vector component of ***w***.

2 Because the eigenvalues of a product of two triangular matrices is equal to the product of their eigenvalues.

1 Note that *N*_*I*_ does not necessarily equal *N*_*E*_.

2 The choice of the L1-norm is motivated by the synaptic competition for a fixed amount of resources, where, in the simplest case, each unit of resource linearly increases synaptic strengths. Higher-order L-norms do not affect the learning of feedforward receptive fields. However, in recurrent networks, they can lead to instabilities (cf. Sec. 4.3).

1 Note that ***A*** and ***VA*** are of dimension *N*_*E*_ × *N*_*I*_, and 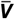 is of dimension (*N*_*E*_ + *N*_*I*_) × (*N*_*E*_ + *N*_*I*_).

2 To show that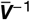 is the inverse of 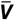it is useful to define the Moore-Penrose inverse A^−1^ = ***A***^⊤^ **(*AA***^⊤^) ^−1^ and note that ***A***^⊤^ (𝟙 − ***AA***^⊤^)^−1^ = (𝟙 − ***AA***^⊤^) ^− 1^***A***^⊤^

3 Similarly, each additional laterally projecting excitatory neuron adds another null eigenvector. In that case, the lateral excitatory weight component and the feedforward excitatory weight component have opposite signs such that they cancel each other (cf. Sec. 5.1).

1 For ***K***_*E*_ = 1 and ***K***_*I*_ = ***AA***^⊤^ −1 columns of ***W***^*^ holds null eigenvectors that can be formed by a linear combination of null eigenvectors 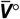 given in Eq. 54.

2 In general, this is not the case for null eigenvectors. Following the same formalism for null eigenvectors 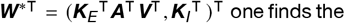 condition that ***A***^⊤^ **Λ*A*** must be diagonal. In general, this is not the case, e.g., when multiple inhibitory neurons are tuned to the same eigenvector, i.e., when multiple columns of ***A*** hold the same vector up to a constant factor.

1 Note that ker ***(A***^⊤^) = ker ***(AA***^⊤^) and ker ***A*** = ker ***(A***^⊤^ ***A)***, for any matrix ***A***.

2 Note that we must make use of the inverse instead of the transpose since, in general, the eigenvector matrix 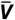 is not orthonormal.

1 Note that when sorted, ***A***^⊤^ ***A*** is a block diagonal matrix. Further, as noted before, the matrix ***AA***T is always diagonal.

2 The dimensionalities of these blocks depend on the number inhibitory neurons tuned to the respective feedforward eigenvector, i.e., if there are 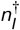 inhibitory neurons tuned to a specific feedforward eigenvector ***v***^†^, the dimensionality of the corresponding block is 1 + *n*_*I*_, due to one regular eigenvector and 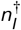 corresponding null eigenvectors.

1 Since we assumed that there is exactly one inhibitory neuron per feedforward eigenvector, there is also exactly one null eigenvector per feedforward eigenvector (cf. Eq. 52)

1 Here, we make use of the equality 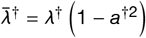.

2 Note that we consider non-repulsive fixed points 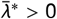 and inhibitory neurons with positive firing rates, i.e., *a* > 0, ∀*a*, such that the second term in the second stability condition is always negative.

1 Here, 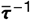 and ***K***^−1/2^ are both diagonal matrices and commute.

1 *a*^*2^ = ***a***^*⊤^ ***a***^*^, cf. Eq. 80.

2 The new modified covariance matrix in the original coordinates is 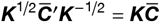, with eigenvectors 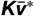.

3 We would have to employ the eigenvector basis of the new modified covariance matrix 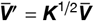 for triangularization.

4 From Eq. 103 we see that we can freely shift scaling matrices ***K*** between the modified covariance matrix and the plasticity timescale by pulling diagonal matrices of the same shape as ***K*** through the bracket (cf. Eq. 102).

5 A change in the overall weight norms, however, can affect the magnitude of postsynaptic activities and synaptic changes.

1 In this case, 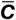 is a true covariance matrix, since the lateral projecting neuron is excitatory.

1 More precisely, ***q*** would converge to a multiple of the principal eigenvector. Here, we consider the more general case where ***q*** is proportional to an arbitrary eigenvector. We will see that with suitable lateral input, any feedforward eigenvector can be stable.

2 However, the constraint term in the weight dynamics introduces interactions between the subspaces orthogonal and parallel to ***q***.

3 Note that ***C***^−1^ = (***C***^−1 ⊤^ since ***C*** is a true covariance matrix, i.e., ***C*** and ***C***^−1^ are symmetric.

1 If *a*_*q*_ is too small so that 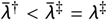, the principal feedforward eigenvector ***v***^‡^ of ***C*** with eigenvalue *λ*^‡^ is stable and 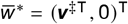.

2 If none of the laterally projecting neurons is tuned to a specific feedforward eigenvector ***v***^‡^, i.e., ***v***^‡^ ⊥ ***q***_*i*_ ∀*i*, the corresponding fixed point becomes 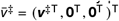.

3 Another example is neurons that project from outside the local circuit, e.g., from another brain area that is higher up in the processing hierarchy.

1 Since we assume Hebbian plasticity between all types of neurons, excitatory and inhibitory, we do not specify the neuron type. *y*_*a*_ and *y*_*b*_ are the firing rates of two arbitrary vectors that are part of two diffe rent eigencircuits.

2 The vectors (***w***^⊤^, ***w***_*E*_^⊤^ ) ^⊤^ and ***w***_*I*_ are normalized such that ∥ (***w***^⊤^, ***w***_*E*_^⊤^ ) ^⊤^ ∥ _*p*_ = *W*_*E*_, and ∥***w***_*I*_ ∥_*p*_ = *W*_*I*_. This is achieved by scaling the excitatory and inhibitory part of the regular eigenvector, i.e., scaling ***v***_*E*_ by *k*_*E*_ = *W*_*E*_ / ∥***v***_*E*_ ∥_*p*_, and ***v***_*I*_ by *k*_*I*_ = *W*_*I*_/ ∥***v***_*I*_ ∥ _*p*_ (cf. Sec. 2.1.3)

1 Note that we allow self-excitation and self-inhibition, i.e., in a fully connected recurrent network, *σ*_*r*_ also appears on the right sides of the equation, as an entry of ***σ***^*E*^ or ***σ***^*I*^.

2 Note that for 0 > Φ < 1 there exists a real solution for *σ*_*E*_, independent of *n*_*E*_.

1 We will show in Section 5.1 that eigencircuits are in fact fixed points of the weight dynamics.

2 See Eq. 146 for the definition of the eigencircuit attraction *λ*_eig_.

1 We omitted the angle notation ⟨· ⟩ to improve readability.

2 It does not affect the sign of the eigenvalues of the Jacobian, since *τ* is always positive. In principle, however, different timescales for excitatory and inhibitory weights can affect stability (cf. Sec. 2.2).

3 Note that here the superscript ‘^*^’ indicates a variable that is evaluated in the fixed point of the weight dynamics and not a fixed point of the firing rate activity. Different input patterns ***y*** result in different neural activities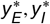.

1 In our simulations, we constrain synaptic weights to be positive. Then null eigenvectors with negative weights are only relevant in combination with regular eigenvectors: When a null eigenvector is added to a regular eigenvector, the net synaptic input does not change. For example, a decrease in recurrent excitation due to a negative excitatory component of the null eigenvector is balanced by an increase in feedforward excitation.

1 We presume that when considering the stability of non-eigenvector fixed points, it is possible to make a similar argument as in Section 2.2.3 and consider regular eigenvectors of a different modified covariance matrix 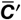 with adjusted plasticity timescales, *k*_*E*_*τ*_*E*_, *k*_*I*_*τ*_*I*_. Here we consider the case of *τ*_*E*_ = *τ*_*I*_ and regular eigenvectors of 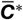, i.e., *k*_*E*_ = *k*_*1*_ = 1.

2 Remember that we set 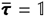.

3 In general, there are 2 × (*n*_*E*_ + *n*_*I*_) normalization terms.

1 Note that in our specific network the top left block of 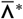 is equal to 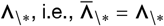, because there are no neurons tuned to the respective feedforward eigenvectors. In particular, 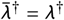 (cf. Eq. 151)

1 This is due to the potentially non-zero elements in the block below the top left diagonal block of 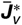 in Equation 195.

1 Note that also the tuning of the inhibitory neuron changes, although it is not directly perturbed.

1 Recall that 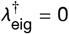 and, therefore, 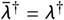 (Eq. 151).

2 The Jacobian of the original circuit (Fig. S3*C*) has additional entries to the right of the top left diagonal block in Equation 195 that are non-zero. These non-zero entries result in the growth of synapses of 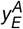 in the direction of eigencircuit *B* due to ‘second-order’ perturbations of recurrent synapses from eigencircuit *A* to eigencircuit *B*.

3 Non-zero entries of the Jacobian to the right of the top left diagonal block are cancelled by the zero entries in the initial perturbation vector 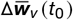 (cf. Eq. 195).

4 Note that the term in Equation 203 that corresponds to the tuning shift of the inhibitory neuron can be positive since d*y*_*I*_ d (***w***_*IF*_T) is negative when the circuit is inhibition stabilized (*6*).

5 See Section 5.2.3 for a discussion of the case *β*_*E*/*I*_ * 0.

6 An unoccupied input mode corresponds to 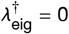 (cf. Eq. 146).

